# Deciphering the structural intricacy in virulence effectors for proton-motive force mediated unfolding and type-III protein secretion

**DOI:** 10.1101/557496

**Authors:** Basavraj Khanppnavar, Anupam Roy, Kousik Chandra, Nakul Chandra Maiti, Saumen Datta

## Abstract

Many gram-negative pathogenic bacteria use type III secretion system (T3SS) to inject virulence effectors directly into the cytosol of targeted host cells. Given that the protein unfolding requisite for secretion via nano-size pore of T3SS injectisome is an energetically unfavorable process, “How do pathogenic bacteria unfold and secrete hundreds of toxic proteins in seconds” remain largely unknown. In this study, first, from an in-depth analysis of folding and stability of T3SS effector ExoY, we show that the proton-concentration gradient (∼pH 5.8-6.0) generated by proton-motive force (PMF) can significantly amortize tertiary structural folding and stability of effectors without significant entropic cost. Strikingly, it was found that the lower energetic cost associated with the global unfolding of ExoY is mainly due to its weakly folded geometry and abundance of geometrical frustrations stemming from buried water molecules and native-like folded intermediates in the folded cores. From in-silico structural analysis of 371 T3SS effectors, it can be curtained that T3SS effectors belong to typical class (disorder globules) of IDPs and have evolved similar conserved intrinsic structural archetypes to mediate early-stage unfolding. The slower folding kinetics in effector proteins requisite for efficient T3SS-mediated secretion mostly stems from reduced hydrophobic density and enhanced polar-polar repulsive interactions in their sequence landscapes. Lastly, the positively evolved histidine-mediated stabilizing interactions and gate-keeper residues in effector proteins shed light on collaborative role of evolved structural chemistry in T3SS effectors and PMF in the spatial-temporal regulation of effector folding and stability essential for maintaining balance in secretion and function trade-off.

## INTRODUCTION

For invading pathogenic bacteria, secretion of virulence effectors into the cytosol of host cell remains a key step in seizing the host cell machinery in their favor (1). To facilitate this, many Gram-negative pathogenic bacteria have evolved type-III secretion system (T3SS) for injecting arsenals of virulence effectors directly into the cytosol of the host cells (2, 3). T3SS is a megadalton proteinaceous nanostructure (also known as injectisome), spanning bacterial and host cell membranes, to form a continuous needle-like conduit through which virulence effectors are secreted (3). Interestingly, the inner diameter of T3SS injectisome is just 25-35 Ǻ (4) and thereby requires at least partial unfolding of effectors to get secreted through a narrow passage (5). Although, in general, T3SS ATPase present in the entry site of injectisome is believed to play a predominant role for effector protein unfolding and T3SS secretion (6, 7). Nevertheless, effector unfolding is a more complicated process as it also relies on several players and factors, including T3SS chaperones, PMF, effector protein stability, and folding kinetics (7–12). Lately, it has been also found that the proton motive force (PMF) is capable of executing effector unfolding and T3SS secretion, in ATPase independent manner (13, 14). Thereby, suggesting that T3SS ATPase is not absolutely required, but rather plays a complementary role in effector unfolding and T3SS secretion.

Among the components of PMF, both electric potential gradient (ΔΨ) and proton concentration gradient (ΔpH) are found to be equally essential for T3SS secretion irrespective of presence or absence of functional ATPase (13–17). Nonetheless, in flagellar T3SS system, an electric potential gradient is only required for effector secretion, but in absence of functional ATPase, both electric potential gradient and proton concentration gradient are essential (14, 15, 18–20). Owning to the fact that the flagellar secretion system is a more specialized system and needs to secrete a limited number of conserved effector proteins for biogenesis of flagella (2, 21–23), whereas, virulence-associated T3SS has to secrete a multitude of diversified effectors (1, 3, 24). Thus, the differential dependency of these secretory systems for proton-concentration gradient seems to correlate with the evolutionary nature of effectors secreted. Concurrently, studies also show that proton concentration gradient of PMF can only generate mildly acidic conditions i.e. around pH 5.8-6.0, which in general is insufficient for acid-unfolding of larger effector proteins (**Fig. S1**), given that protein unfolding is an energetically unfavorable process (16). Thus, in order to shed mechanistic insights on PMF-mediated effector unfolding and secretion, it is necessary to explore pH-dependent folding and stability of T3SS effectors.

Contemporary studies also show that T3SS mediated secretion requires distinct kind of substrates for efficient secretion. As very stable proteins such as green fluorescence protein (GFP), glutathione S-transferase (GST), Ubiquitin, or rapidly folding proteins such DHFR when fused to T3SS secretions signal, remain incompatible for T3SS secretion (5, 9–11). However, upon engineering these fusion proteins with mutations that destabilize them, the blockade in T3SS secretion vanishes completely (9). Thus, suggesting that the folding and stability of T3SS effectors is possibly being modulated for rendering effector unfolding and T3SS secretion. Nevertheless, considering the fact that the number of unique effectors secreted into the host cell is comparably less. Instability of an effector may also hamper its ability to function inside the host cell. In this context, “How does the evolution of T3SS effectors negotiate stability and function trade-off? What are deterministic factors evolved by effectors for slowing down its folding kinetics? To address these questions, and to gain mechanistic insights on T3SS secretion, a comprehensive investigation of physiochemical and structural properties of T3SS effectors is warranted.

In the current investigation, systematic efforts combining experiments and computations have been employed to shade mechanistic insight of effector unfolding and T3SS secretion. First, the pH-dependent folding and stability of ExoY, a 43kDa T3SS effector from *Pseudomonas aeruginosa*(25) was explored using NMR, Raman, FT-IR, and other biophysical techniques. Further, to scrutinize our findings and its general implications, statistical and computational analysis of structural and physiochemical properties of other T3SS effectors was also carried out. In sum, our results provide new fundamental insights on molecular mechanism evolved by Gram-negative pathogenic bacteria to secrete virulence effectors or toxins into the targeted host cells.

## RESULTS AND DISCUSSION

### Role of the proton-concentration gradient of PMF in effector unfolding and T3SS secretion

Contemporary studies suggest that the proton-concentration gradient (ΔpH) generated by the proton-motive force (PMF) play a vital role in effector unfolding and T3SS secretion (13–15, 18). However, given that protein unfolding is an entropically unfavorable process and PMF can only generate mild acidic conditions (∼ pH 5.8-6.0) (15, 17, 20), the role of PMF in effector unfolding still remains an enigma. To understand this at the molecular level, pH-dependent folding and stability of ExoY, a T3SS effector from *P. aeruginosa* was investigated. This 43 kDa effector protein is sufficiently larger in size to represent averaged sized effector proteins (**Fig. S1**) and can be characterized using multiple biophysical techniques. More, it also lacks secretion chaperone, and thereby, it is also a useful model system to gain a detailed understanding of a chaperone-independent unfolding and T3SS secretion. From CD spectroscopic analyses, it was observed that a mild acidic condition of PMF could only trigger a minor loss in secondary structural contents of ExoY (Fig. 1A). However, with decreases in pH from 7.4 to 5.8, the melting temperature (*T*_m_) of ExoY sharply decreased from 42.7 °C to 37.9 °C, which is close to the ambient physiological temperature of eukaryotic hosts (Fig. 1B & C). Thus, suggesting that a mild acidic condition of PMF, even with the partial unfolding of secondary structures, can significantly reduce the overall structural stability of ExoY.

**Figure 1.**
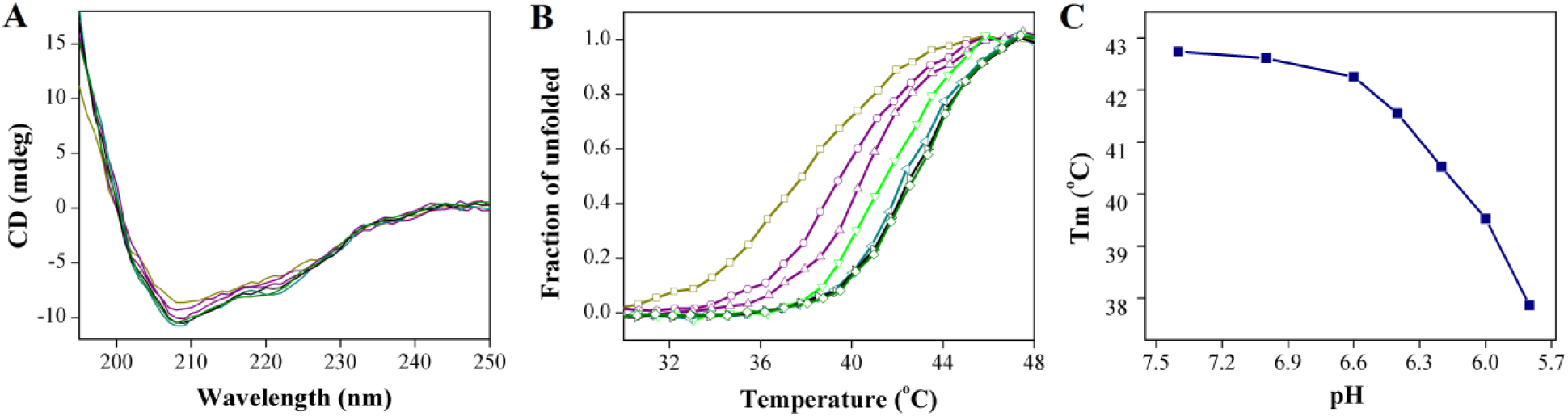
Folding and stability of ExoY in presence of mild acidic conditions of proton-motive force (PMF). **(A)** Far UV CD spectroscopy analysis showing a trivial loss in ordered secondary structural contents of ExoY with lowering of pH from 7.4 to 5.8. **(B)** However, the thermal denaturation studies showed a more significant loss in the overall stability of ExoY as perceptible from melting temperature (*T*_m_) of each sigmoidal thermal unfolding profile. The *T*_m_ of ExoY at pH 7.4 (Olive), pH 7.0 (Black), pH 6.6 (Dark Cyan), pH 6.4 (Green), pH 6.2 (Violet), pH 6.0 (Purple) and pH 5.8 (Dark Yellow) were 42.74 °C, 42.6 °C, 42.3 °C, 41.6 °C, 40.5 °C, 39.5 °C, 37.9 °C respectively. The thermal melting of ExoY was monitored at 222 nm. **(C)** From the plot of melting temperature versus pH, it can be clearly seen that the thermal stability of ExoY decreases more drastically after pH 6.4 onwards.

To further delineate the pH-dependent conformational transitions, vibrational spectroscopic analysis of ExoY was carried out. Off-resonance Raman spectrum of ExoY crystals showed that the native state (at pH 7.4) is predominated by a broad amide I band with a center of 1657 cm^−1^ and bandwidth at half height (BWHH) of ∼44 cm^−1^ (Fig. 2A). This mode of vibrations corroborates the existence of structural heterogeneity with a significant content of molten unordered hydrated conformers (vibrational band at 1645 cm^−1^) and extended PPII structures (vibrational band at 1681 cm^−1^) along with predominant helical and beta conformers at 1657 cm^−1^ and 1671 cm^−1^ respectively (26–28) (Fig. 2A, **Table S1**). With a gradual exposure of ExoY crystals to pH 5.8, the amide I band broadened (BWHH increased from 44 cm^−1^ to 49 cm-^1^), suggesting an additional increment in structural heterogeneity (Fig. 2B & C). In addition, the major amide I vibrational band at 1657 cm^−1^ for α-helices was significantly reduced, whereas vibrational bands associated with extended β-strand (1670-1675 cm^−1^) and poly-l-proline II (PPII) conformations (1681-1687 cm^−1^) were found to be more prominent at pH 5.8 (Fig. 2, **Table S1**).

**Figure 2.**
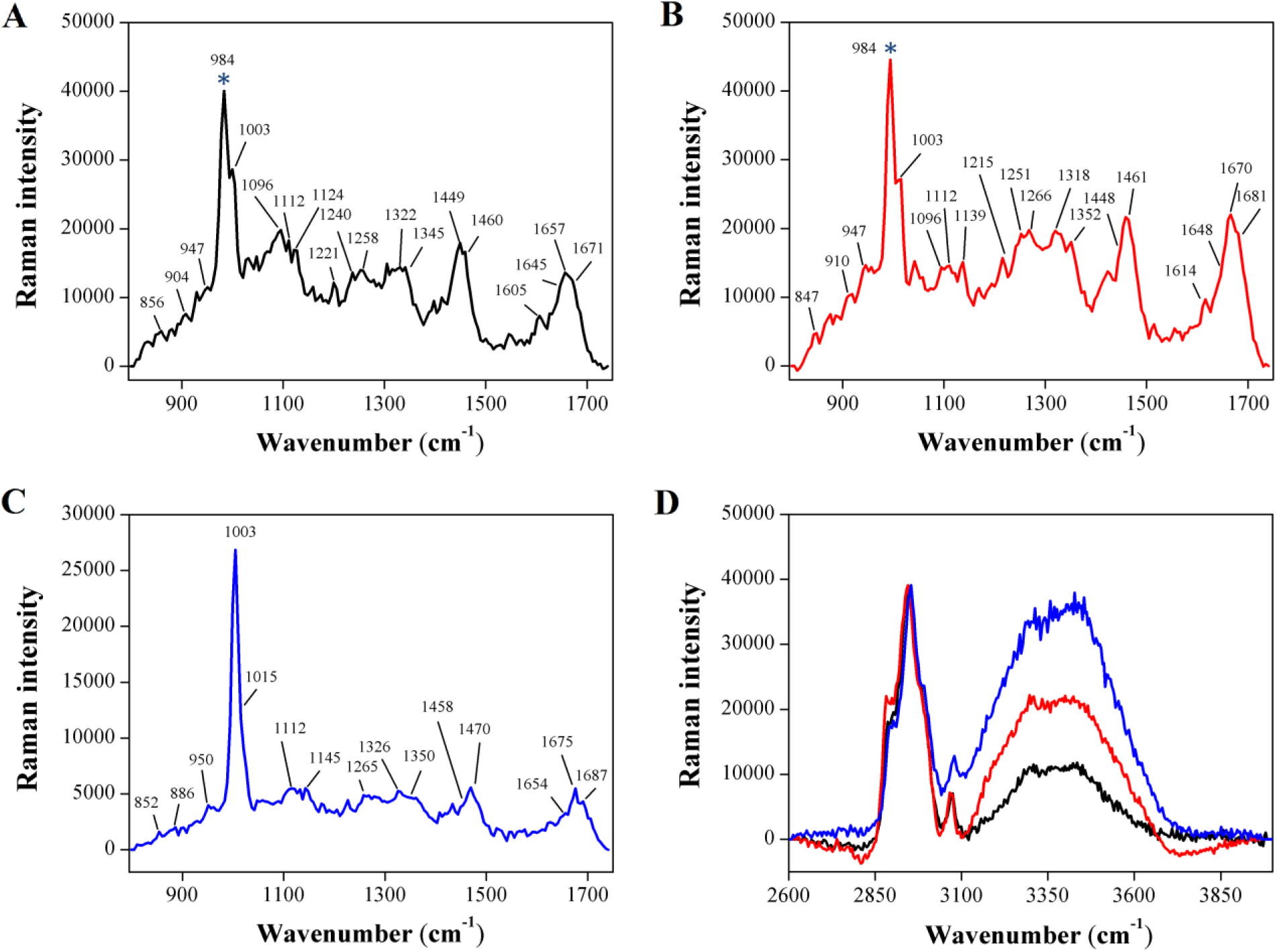
Raman crystallographic analyses of ExoY single crystals. 633 nm He-Ne laser excited Raman spectra (800-1800 cm^−1^) of ExoY crystals at pH 7.4 **(A)**, and at pH 5.8 after 30 minutes **(B)** and 60 minutes **(C)** incubation period. Spectral assignments of some of the characteristic Raman vibrational bands from the fingerprint spectral region (800 - 1800 cm^−1^) are tabulated in **Table S1**. **(D)** Raman spectra of ExoY in the frequency range of CH and OH stretching vibrations obtained from protein crystal at pH 7.4 (black line), and at pH 5.8 after 30 minutes (red line) and 60 minutes (blue line) incubation period. The sharp peak at 981 cm^−1^ (indicated by *) was due to P=O bending mode (symmetric) of phosphate ion embedded with a single crystal of ExoY.

A similar trend in the secondary structural unfolding of ExoY was also observed from FT-IR spectroscopic studies (**Fig. S2A, Table S2**). The FT-IR amide I band of ExoY at pH 7.4 primarily consisted of a broad amide I spectrum (1640-1648 cm^−1^) mainly due to the presence of hydrated molten helical conformers or (3/10)-helix conformers (29, 30) along with sharp amide I vibrational band (1653 cm^−1^) specific for pure α-helical conformations. In addition, the amide I spectrum of ExoY also showed weak vibrational bands specific for extended PPII structure (∼ 1633 cm^−1^), extended β-strand conformers (1625 cm^−1^) and β-turn marker bands at 1662 cm^−1^ and 1679 cm^−1^, respectively (29, 31). With decrease in pH to 5.8, the intensity of amide I bands for extended PPII conformation (1631 cm^−1^) and extended β-strand conformers (1626 cm^−1^) were found to be elevated, whereas the intensity of amide I bands specific for pure α-helical conformers (1650-1654 cm^−1^) and intermediate helical conformations (1639-1648 cm^−1^) was significantly reduced (**Fig. S2B** & **Table S2**). Moreover, a simultaneous increase in β-turn signature could also be visualized from sharpened vibrational signatory bands at 1660 and 1695 cm^−1^. These significant changes can be more clearly visualized by comparing decreased helical signatures with increased signatures of extended PPII and β-turn structures. Thus, in sum, these observations clearly demonstrate that the minor loss in secondary structural contents of ExoY observed in CD spectra and increased pool of PPII and random β-turn conformers at pH 5.8 is mainly due to conversion of intermediate hydrated molten helical conformers present in native structure and to some extent from pure α-helical structures (Fig. 1A, Fig. 2 & **Fig. S2**).

From the analysis of tertiary structural signatures, the intensity of vibrational bands (1449 cm^−1^) specific for total CH-, CH_2_-, CH_3_-deformation and scissoring motions at pH 5.8, with respect to marker band (Phe) at 1003 cm^−1^, was significantly reduced (Fig. 2). Similarly, the non-amide Raman signatory bands (1000-1200 cm^−1^) encoding overall backbone and side chain information (26, 27), were also distorted markedly, suggesting a significant loss in overall tertiary structural integrity of ExoY. Intrinsic tryptophan fluorescence analysis also suggested the considerable loss in its tertiary structural integrity of ExoY with gradual lowering of pH (**Fig. S3A & B**). Furthermore, as protein unfolding is primarily characterized by increased backbone solvation, the increased intensity of Raman OH stretching bands (3100-3800 cm^−1^) at pH 5.8, clearly depict the increased backbone solvation and partial unfolding of ExoY (Fig. 2D). From hydrogen-deuterium (H/D) amide exchange kinetic experiments, it can be clearly seen that with a gradual lowering of pH 7.4 to 5.8, there is a sharp increase in the amide exchange rate of ExoY **(Fig. S2C, D & E)**. Thus, in sum, these observations clearly demonstrate that the mildly acidic conditions of PMF can significantly tweak tertiary structural intricacy of ExoY to facilitate its unfolding and secretion via nano-size pore of T3SS injectisome.

Even though protein unfolding is energetically unfavorable, yet a significant loss in the overall tertiary structure of ExoY was observed in presence of mildly acidic conditions of PMF. To understand the underlying factors that are responsible for such remarkably unfavorable process, thermodynamic parameters for PMF-mediated acid unfolding of ExoY were probed using isothermal calorimetric (ITC) titration experiments. **Figure S4** shows a typical ITC titration curve obtained after titrating ExoY with acid (from pH 7.8 to 5.6). The overall enthalpic contribution in PMF-mediated unfolding of ExoY was found to be thermodynamically favorable (Δ*H°*= −6.4 Kcal/mol) compared to its entropic feasibility (Δ*S°*= 1.8 cal/mol). The small positive entropic cost associated with the unfolding of ExoY suggested that the overall exposure of buried hydrophobic surface is apparently smaller than seen in the general unfolding of globular proteins. Consistent with this observation, in hydrophobic dye binding assay, the exposed hydrophobic surfaces of ExoY at pH 5.8 was very much trivial (only 3.2%) compared to its completely unfolded state at pH 2.0 (**Fig. S3C, Fig S5B & C**). Interestingly, the major shift in the exposure of hydrophobic surface apparently occurs after the ambient pH of PMF. Furthermore, due to significant exposure of hydrophobic surfaces below pH 5.8, ExoY also showed misfolding propensity during CD melting experiments (**Fig. S5D**). Therefore, it can be regarded that PMF-mediated unfolding of effectors is a more evolved phenomenon that mainly involves exposure of the polar tracts of tertiary structures than non-polar or hydrophobic residues. From the significantly decreased stability of ExoY with negative enthalpic changes, it seems that the energy required for disrupting the overall tertiary structural contacts is relatively less. This can be essentially true when there are conserved solvent-exposed large polar/charged residues inside the folded cores and there is a restriction in non-covalent interactions (i.e. hydrogen bonding and salt bridge interactions) essential for maintaining the labile partially folded tertiary structure of ExoY.

### Geometrical frustration in folded core minimize the energetic cost of the global unfolding of ExoY

Protein folding is primarily an entropically driven process where the enthalpic cost of unfavorable hydrophobic-solvent contacts is minimized by removing bulk solvent or water molecules (32, 33). In the early stages of protein folding, the non-polar side chains of hydrophobic residues undergo hydrophobic collapse and peptide backbones form ordered secondary structures, only to circumvent water molecules (33–35). Likewise, inter-molecular tertiary contacts or interactions at a later stage of protein folding is also critical for removing residual water molecules and to attain more compactly folded geometry that can prevent subsequent water penetration inside the folded core of the protein (32, 36, 37). In this context, a degree of protein folding and stability can be easily estimated from its hydrophobic collapse or packing, ordered secondary structural content, the degree of co-existence of secondary structures in hydrophobic packing, overall tertiary structural contacts, and level of water solvation. Based on the following parameters, a protein with highest folded geometry is relatively more stable and can resist its unfolding in presence of different external stimulates.

To realize the interplay of the intermolecular forces and the molecular basis of susceptibility of ExoY to mildly acidic conditions of PMF, we analyzed the core geometrical and structural features embedded in the native crystal structure of ExoY. Interestingly, even though the crystal structure of ExoY was obtained after removing large proportions of unordered structures (38) (∼ 28.6% residues), still, in the crystal structure, almost 45.4 percent of residues encoded less ordered or extended/coil conformations and remaining 54.6 percent of residues were ordered secondary structures i.e. helical and beta structures. More interestingly, the majority of extended/coil conformations in crystal structure exist as disordered-globule inside overall folded geometry of ExoY (Fig. 3A). As disordered globules are geometrically frustrated native-like folded intermediates, the existence of such disordered-globules in the heart of folded core not only form a barrier between folded sub-domains but will also compromise the overall tertiary structural stability of ExoY. From the analysis of hydrophobic packing and dispersion in the crystal structure, it was found that majority of hydrophobic residues or clusters are more dispersed in toroidal shape geometry and do not form the hydrophobic center of masses, as typically seen in globular proteins (Fig. 3B). Interestingly, it was also observed that the domain 1 which encompasses more ordered secondary structural elements has less compact hydrophobic packing i.e. more dispersed hydrophobic residues or clusters in the toroidal shape, whereas the other domain comprising disordered globule and with least ordered secondary structures has maintained relatively more compact hydrophobic packing. Thus, in compliance with these observations, it seems that co-evolution of abundant folded secondary structures and compact hydrophobic packing is avoided in ExoY to render its unfolding energetically less unfeasible.

**Figure 3.**
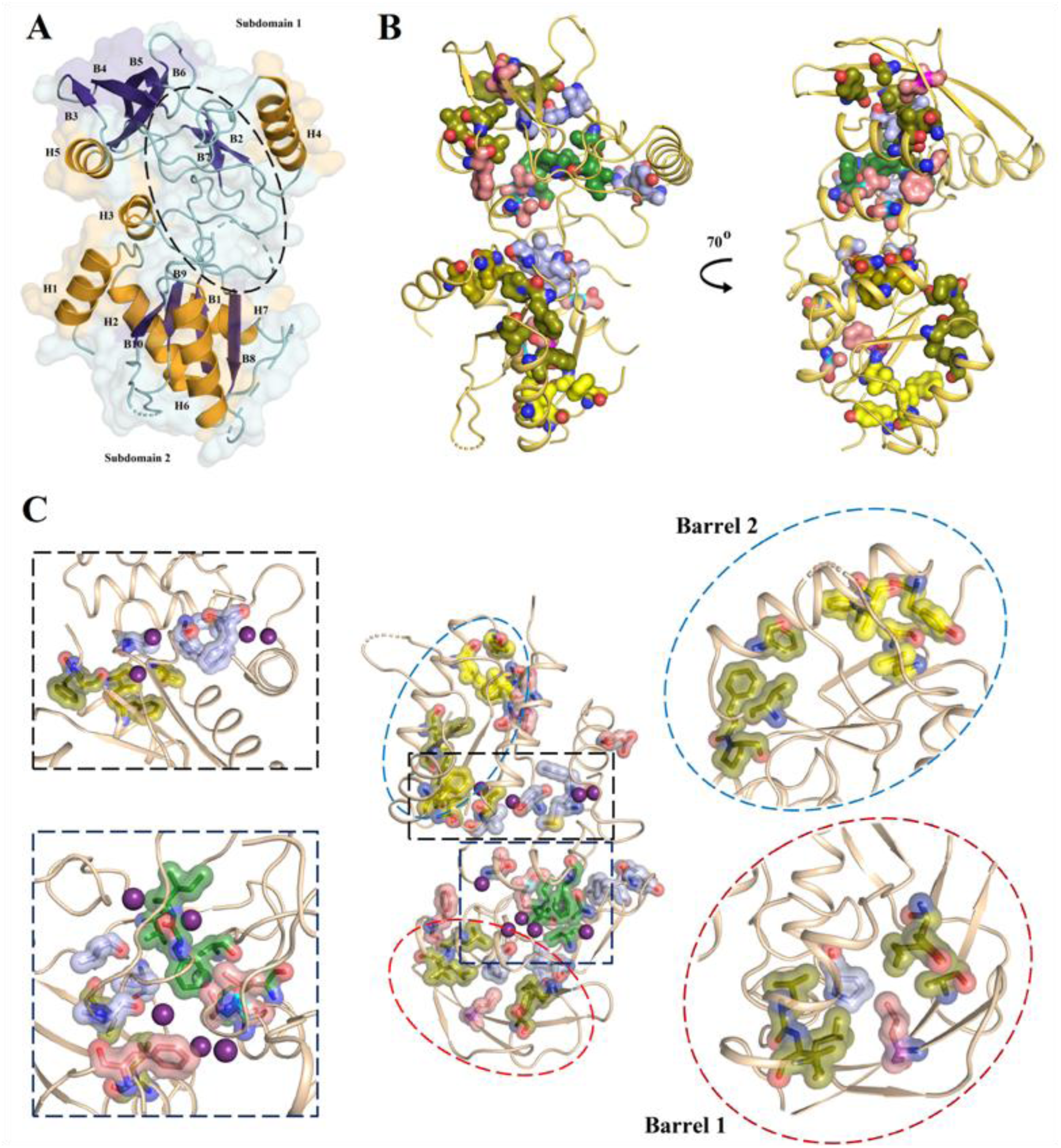
The degree of independent folding and stability in the crystal structure of ExoY. **(A)** The overall bi-domain structure of ExoY (PDB: 5XNW) comprised of two folded globule (subdomains) separated by one disordered globule (highlighted in a dashed circle). **(B)** In both domains of ExoY, the hydrophobic residues were dispersed in the planar toroidal ring-like geometry. Interestingly, the hydrophobic packing in domain 1 (which embedded a larger portion of disordered globule) is relatively more condensed compared to domain 2. Approximately 60-70 degree planar tilting is observed between these two planar hydrophobic toroidal rings. **(C)** Geometrical frustration and comparative folding in the crystal structure of ExoY. The hydrophobic clusters and residues are shown in different color code according to its hydrophobicity strength and its distribution in structure (for more details see **Fig. S9**). Out of eight hydrophobic clusters present in crystal structure of ExoY, only one hard (cluster A) and 3 soft (cluster B, F & G) hydrophobic clusters were seen in relatively stable barrel-like structures and remaining hydrophobic clusters were either distributed in disorder-globule (cluster D, E, & H) or in isolated super-secondary structures i.e. helix-turn-helix (cluster C). In ordered barrel-like structures, each hydrophobic cluster essentially acted as lids of barrel structure made up of one helix and four beta strands. These two stable barrel-like structures were also the only dehydrated regions in the crystal structure of ExoY. Presence of such entrapped buried water molecules inside hydrophobic clusters not only reduces the overall stability of hydrophobic clusters and its connection with neighboring hydrophobic clusters or residues but will also elevate global geometrical frustrations (for more details see **Fig. S10**).

Further, from analysis of hydrophobic clustering strengths and its distribution(34, 36, 39, 40), it was observed that the overall structure of ExoY comprises of 8 hydrophobic clusters, of which only 2 are hard hydrophobic clusters (cluster A & D) and remaining 6 are soft hydrophobic clusters (**Fig. S9**). Interestingly, out of these eight hydrophobic clusters, only one hard (cluster A) and 3 soft (cluster B, F & G) hydrophobic clusters were located in relatively stable barrel-like structures (Barrel 1 and Barrel 2) and the remaining hydrophobic clusters were distributed in disordered globule (cluster D, E, & H) or in isolated super-secondary structures i.e. helix-turn-helix (cluster C) (Fig. 3D, Fig. 4A, **Fig. S9**). Barrel 2 encompasses four beta strands (β1, β8, β9, β10) and one helix (α6), which organized to form barrel-like structure and the hydrophobic cluster A and cluster B forming lids of this empty barrel. Similarly, barrel 1 comprising of four beta strands (β3, β4, β5, β6) and a helix (α5) formed twisted barrel-like structural fold with hydrophobic cluster F and G acting as its lids. However, unlike barrel 2, barrel 1 had two isolated hydrophobic residues (P94 and L164) forming weak connections between its hydrophobic lids (**Fig. S10**). The hydrophobic clusters in disordered globule, despite being in energetically frustrated state, were not found to collapse with neighboring hydrophobic cluster, as the radius of these hydrophobic clusters is either less than critical radius required for collapse or suffered conformational strain due to water solvation (reducing hydrophobic strength)(39–41) (Fig. 3D, **Fig. S10**). As a consequence of this, the hydrophobic clusters in disordered globule are entrapped in geometrically frustrated states and significantly compromise the overall folding and stability of ExoY.

**Figure 4.**
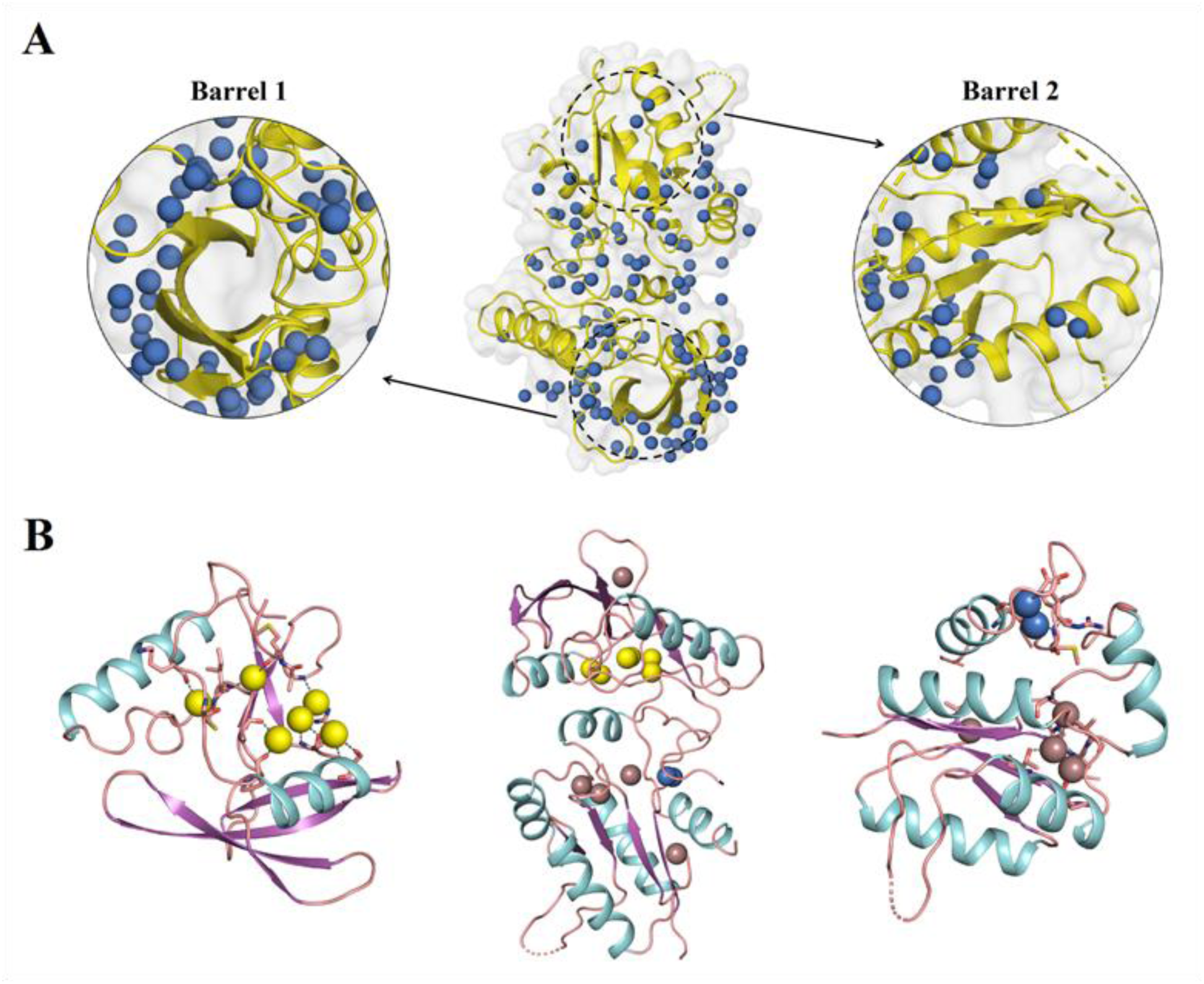
Role of buried water or solvent molecules in the structural stability of ExoY. **(A)** Solvent structure in the native crystal structure of ExoY. Out of 125 water molecules found in the crystal structure, 12 water molecules were found to be buried inside the protein structure, and 116 water molecules were present in the inner hydration shell. Two dehydrated regions in ExoY are highlighted in dashed circles. **(B)** Water molecules inside the disorder globule domain are shown in yellow color. Water molecules in close association with β-strands are shown in grey color, whereas water molecules interacting with helices are shown in blue color respectively. Interestingly, most of the water molecules were found interacting with backbone amide groups. Interaction of water molecules (yellow) with backbone amide bonds (S140, R138, P94, A96 & M209) of loops proceeding β-strands, may possibly have prevented further folding or extension of β-strands. The water molecules which are encapsulated in type-II β-turns/loops and form interaction with backbone atoms may not necessarily have a destabilizing role. As type II β-turn or loops are more ordered structure, here, water molecules may elicit stabilizing effect rather than destabilizing effect. The interaction of backbone atoms of the α7 helix with water molecules (blue) possibly weakens its subsequent backbone interaction propensity crucial for helical geometry and thereby resulting in helix destabilization.

From the analysis of tertiary contacts, surprisingly, even though ExoY contained a large number of polar and charged residues, only 7 charged residues were found to form long-range electrostatic interactions (**Fig. S12A**). Remaining charged residues were either found to be solvated with surface bulk water molecules or formed water-mediated dipolar interactions with side chains of other residues (**Fig. S12B**). More, due to lack of compact hydrophobic center of masses, lower abundance of ordered secondary structure and existences of few long-range tertiary contacts, the overall tertiary structural geometry of ExoY was less compactly folded and lots of bulk water/solvent molecules were found to penetrate the native folded tertiary structure (except two folded barrel-like geometry) (Fig 3D, Fig. 4A and **Fig. S10**). Furthermore, the bulk water molecules were either found to be involved in secondary structural distortions (**Fig. S14B**) or stabilized the molten conformers such as coils, type-II β turn, and loops in disordered globule to prevent further secondary structural folding (Fig. 4B). Thus, the interactions of water molecules seem to accomplish multiple roles, right from destabilization of hydrophobic clusters and isolated helices to stabilization of molten secondary structural conformers.

Given to the fact that protein folding is a process of de-solvation of backbone and side chains, the reverse holds true for protein unfolding, making it a process of solvation of overall tertiary structural geometry (33, 36, 37). From preexistence of partially solvated and intermediately folded structures especially inside the folded core, it seems that ExoY is kind of an intermediately folded protein. Due to such intermediately folded and geometrically frustrated structural features, the slightest perturbation in solvent dynamics by increasing temperature or ionic strength can significantly elevate water solvation in disordered globule, eventually melting disordered globule to form more extended and largely backbone solvated PPII-like structures. This will also lead to the separation of two sub-domains. From the decoding of structural stability of relatively more folded sub-domain 2, it can be observed that interaction of folded barrel 2 with interfacial helices is mostly governed by few weak hydrophobic interactions (**Fig. S11**). These crucial hydrophobic zipping interactions were further weakened by the presence of small hydrophobic (such as alanine) and amphipathic amino acids (such as methionine) or by the existence of bulk water molecules at the interfaces of helices and barrel-like structures.

Interestingly, other than these weak hydrophobic interactions, only a few histidine-mediated hydrogen bonding interactions were found to stabilize the interaction between isolated helices with barrel-like structures and/or with disordered globule (Fig. 7, **Fig. S11**). Given the background that the side chains of histidine residues show pKa value of ∼6.2 - 6.5, these histidine mediated interactions are likely to be disrupted in presence of mild acidic conditions of PMF. As a result of which, zipping interactions between barrel-2 structures and interfacial amphipathic helices of sub-domain 2 will be lost, eventually destabilizing isolated helices. Consistently, in presence of mild acidic conditions of PMF, a significant loss in molten helical conformers and increase in PPII structures, along with dramatic alteration in hydration level (backbone solvation), was clearly evident in the secondary and tertiary structural unfolding studies of ExoY (Fig. 2 **& S2, Table S1 & S2**). Owning to the fact that exposed flexible and molten helical structures had been removed in ExoY crystals, the increased backbone water solvation and poly-proline structures clearly depicted the partial unfolding of disordered globule (Fig. 2). A partial loss in pure α-helical signatures in Raman and FT-IR spectrum, besides conversion of molten helical conformers into extended PPII structures, also clearly suggested the destabilization of helices (Fig. 2 **& S2, Table S1 & S2**). More, as partially unfolded ExoY can behave like a typical block copolymer with two folded barrels as hydrophobic moiety and extended disordered globule comprising coil-coil structures or ELP-like peptide sequences as hydrophilic moiety (42–48) (Fig. 5). In presence of mild acidic conditions of PMF, the increased propensity of ExoY to form coacervated species also clearly suggested the melting of disordered globule and separation of folded sub-domains (Fig. 5, **Fig. S6**). Based on PMF-mediated global tertiary structural unfolding and typical block copolymer induced liquid-liquid phase separation phenomenon, a mechanistic model for effector unfolding and T3SS secretion, and a hypothetical model of partially unfolded ExoY is depicted in **Fig. S16.**

**Figure 5.**
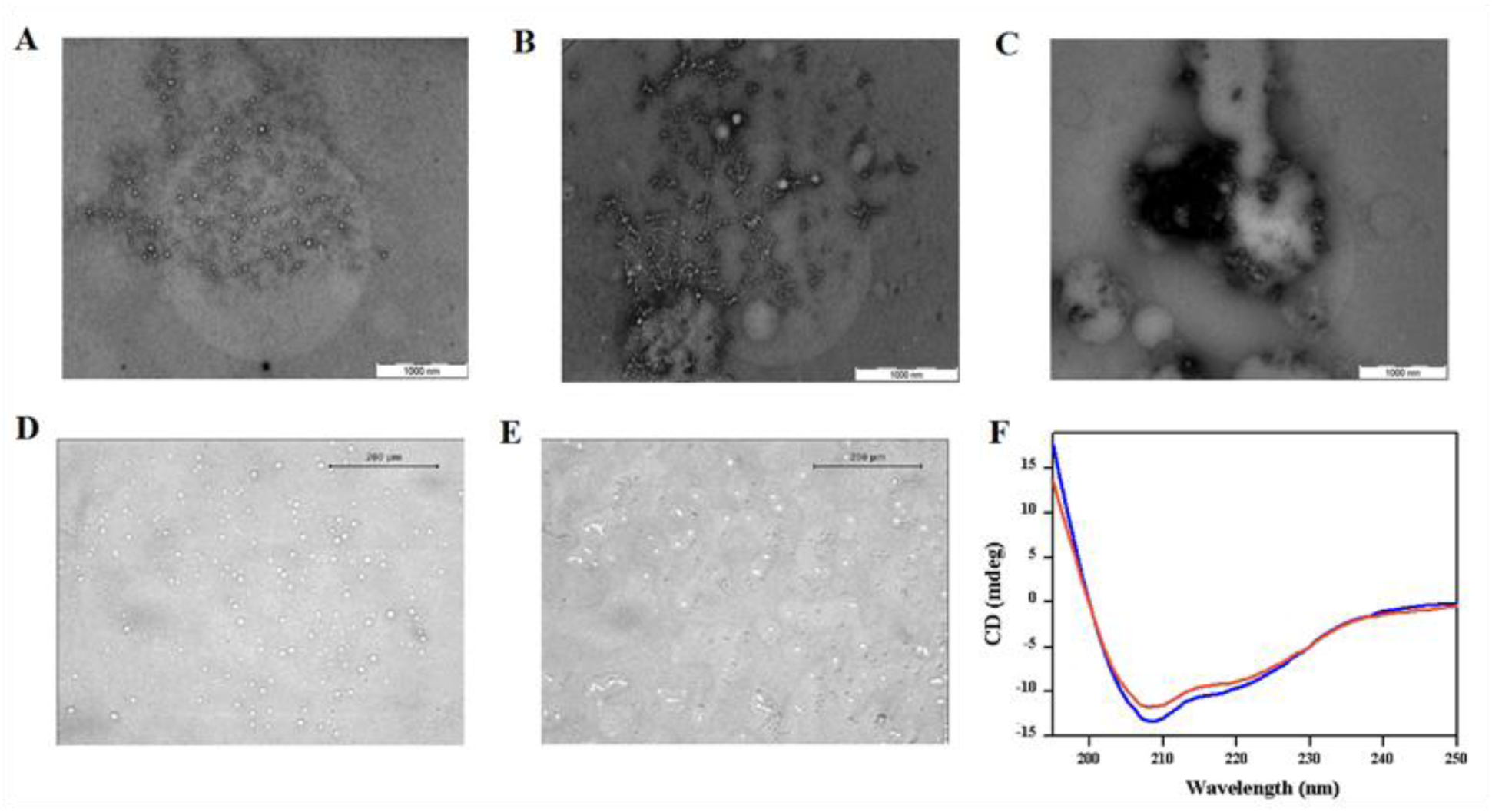
Characterization of IDP-enriched block co-polymeric intermediates of ExoY. Thermo-responsive large scale self-association of ExoY in liquid-liquid phase separated coacervated species, ranging from nanometer to micrometer in diameter as observed from **(A, B, C)** transmission electron microscopic (TEM) and **(D & E)** optical phase contrast microscopic images. **(F)** Far-UV spectrum of ExoY at 15°C, 25°C and 35°C respectively. Thermo-responsive self-association of ExoY does not require significant secondary structural rearrangement or more precisely formation of cross-β structures as observed for typical amyloid-like aggregation (53). Thermo-responsive self-association or liquid-liquid phase separation was mainly triggered by melting of disordered globule (rich in elastin-like peptide sequences) and separation of folded subdomains. As partially unfolded ExoY possesses characteristic features of a typical block copolymer with exposed elastin-like peptide sequences or coil-coil structures of extended disordered globule acting as a hydrophilic moiety and the folded subdomains forming hydrophobic moiety (42–48)

### Negative evolution of folding and stability in effectors to facilitate T3SS secretion

From above structural studies, it can be realized that weak hydrophobic packing, labile tertiary contacts, and increased backbone solvation in the overall structure of ExoY render its unfolding energetically less unfavorable. To scrutinize the existence of similar structural features in other T3SS effectors, we carried out the statistical analysis of amino acid composition in T3SS effectors. From our analysis, it can be clearly observed that the relative frequencies of hydrophobic amino acids (V, L, I, M, W, Y, F) have considerably reduced in T3SS effectors, whereas the frequencies of polar amino acids have increased significantly (Fig. 6A & B). Interestingly, among the hydrophobic amino acids, the frequency of tryptophan, which is the most hydrophobic amino acid and is known for playing a more critical role in early protein folding and nucleation events, was found to decrease more significantly than other hydrophobic amino acids (34, 35). Likewise, other hydrophobic amino acids were also found to decrease according to their hydrophobicity strength and overall effect on protein folding and stability. For instance, the frequency of alanine, which is the smallest and weakest hydrophobic amino acid, was not found to decrease more significantly in T3SS effectors. Similarly, methionine, which is relatively more amphipathic in nature and has a lesser tenacity to participate in hydrophobic zipping interactions, was found to decrease less significantly than all other hydrophobic amino acids (Fig. 6A & B). These observations, in compliance with previous structural analysis of ExoY, clearly depict the existence of weaker overall hydrophobic packing in three-dimensional structures of other T3SS effectors. Furthermore, due to the presence of relatively weaker or amphipathic hydrophobic amino acids, and abundant polar tracts; the strength of existing hydrophobic interactions and clusters will be significantly compromised in T3SS effectors. Consistently, in the crystal structure of ExoY, weak and amphipathic hydrophobic residues (A, Y & M) were the most common interacting residues between stable barrel-like structures and isolated helices (**Fig. S11**).

**Figure 6.**
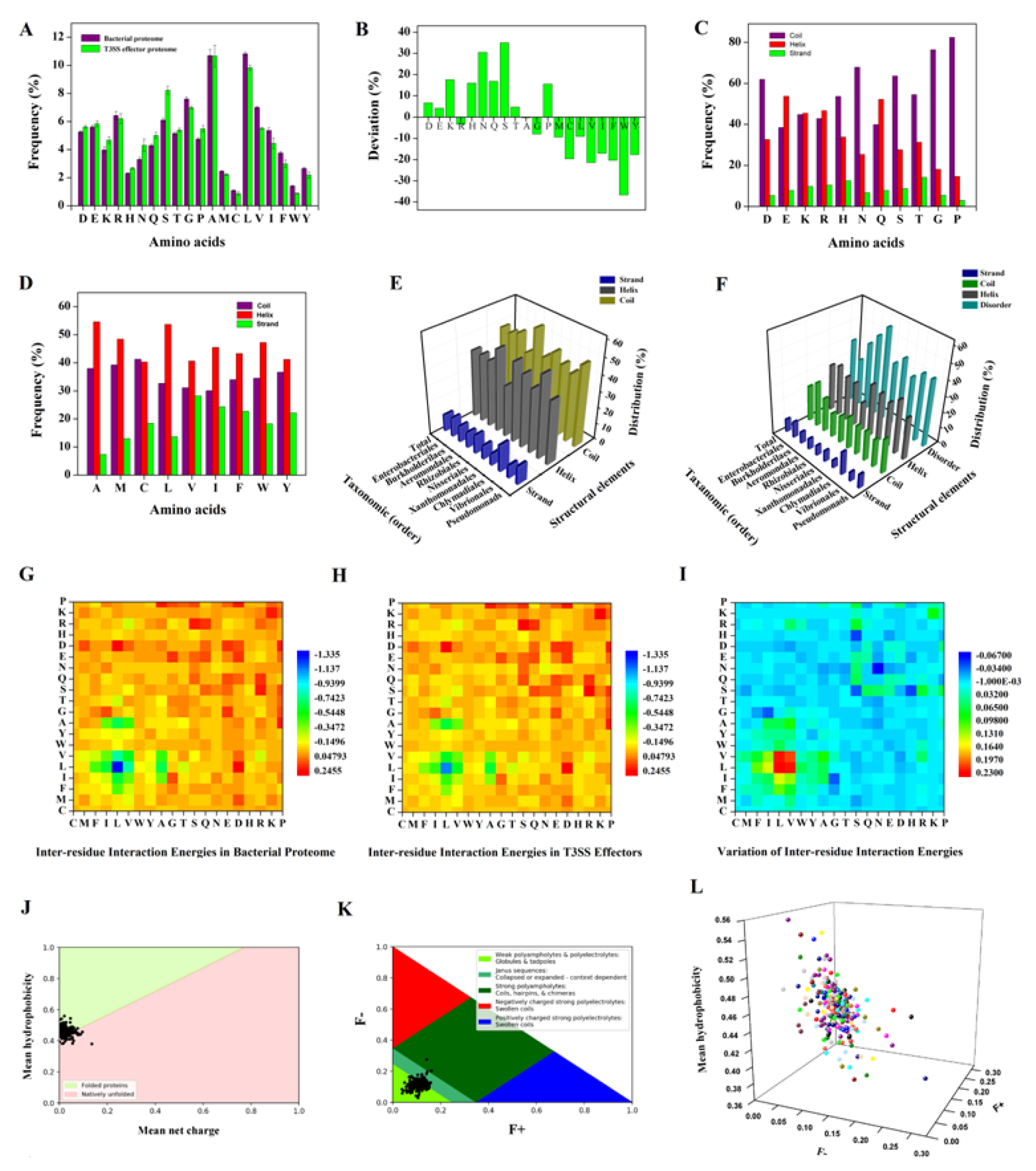
Statistical analyses of physiochemical properties shed light on conserved structural archetypes in effector proteins requisite for T3SS-mediated secretion. **(A)** The average natural occurrence frequency of amino acids in T3SS effectors proteome (Green) and whole bacterial proteome (Purple) of T3SS harboring bacterial species. The natural occurrence frequencies of amino acids in bacterial genomes were obtained from Brbic et al 2016(63) (for complete lists see **Supporting Data 1**). Error bars denote one standard deviation fluctuations. **(B)** The differences in average amino acid content between bacterial proteome and T3SS effector proteome. **(C and D)** Distribution of polar & charged and non-polar amino acids in different secondary structural components of T3SS effectors. Secondary structural architecture in T3SS fitted in **(E)** three components and **(F)** four components showed a rich abundance of flexible and semi-structural elements in T3SS effectors (for complete lists see **Table S3**). **(G)** The inter residue-residue interaction potentials in T3SS effectors, **(H)** control bacterial proteome and **(I)** difference in inter residue-residue interaction potentials between T3SS effectors and control bacterial proteome. The residue-residue interaction potentials between any two types of amino acids were computed based on the probability of interactions of amino acids in the sequence. Standard inter-residue contact potentials were obtained from Thomas et al 1996 (65). Distribution of T3SS effectors in protein folding phase space diagram of **(J)** Uversky plot (55) i.e. mean net charge v/s mean hydrophobicity **(K)** Das-Pappu phase diagram (56) (Fraction of positively charged residues (F+) v/s Fraction of negatively charged residue (F-) **(L)** Three-dimensional charged-hydrophobicity phase diagram (57) (Fraction of positively charged residues (F+) v/s Fraction of negatively charged residue (F-) v/s mean hydrophobicity.

Interestingly, among polar amino acids, the frequencies of polar uncharged amino acids (N, S, Q, and T) were found to increase more significantly than polar charged amino acids (R, D, and E) and the frequency of strong polar amino acids were found to increase more significantly than relatively less polar amino acids (Fig. 6A & B). For instance, the frequencies of serine and asparagine residues have increased more significantly than threonine and glutamine, respectively. Since, most of the highly polar amino acids especially low complexity region (LCR) promoting residues (such as S, Q, and T) and charged residues (such as D, E and K) exhibit higher propensity for unordered or molten secondary structural conformers (49, 50). As a consequence of the positive evolution of these residues along with helix and β-sheet breaker proline residues (51), nearly 39 percentages of residues in T3SS effectors were found to be disordered (**Table S3, Supporting Data 1**). Moreover, as evident from our partially assigned NMR data of ExoY, the higher percentage of disordered residues in effector proteins is not only due to intrinsically unstructured N-terminus secretion signal, but, rather due to the global dispersion of disordered residues in three-dimensional structures of effector proteins (**Fig. S7 & S8**). It can also be observed that nearly 15-20 percentage of NMR predicted disordered residues partially overlapped with disordered loops/regions present in the crystal structure of ExoY (**Fig. S7**). NMR signal for disordered residues, especially, which are in association with hydrophobic collapse or disordered globule, could not be detected, possibly due to large deviation in the longitudinal relaxation time.

Along with the reduced content of ordered helical and β-structures (Fig. 6E, **Table S3**), it can also be seen that both helix and β-structures in T3SS effectors are more amphipathic in nature (Fig. 6C & D) and enclose additional structural stabilizing and destabilizing features (**Fig. S13A, S14 & S15**). Furthermore, from comparison of secondary structure and disorder propensities (see materials and methods section), it was also observed that the nearly ∼14% of predicated ordered residues are very close to the order/disorder boundary, and average percentage of stable ordered secondary structures in effectors is limited to ∼38% only (Fig. 6F, **Table S3**). The decreased ∼14% of ordered structures can be considered as ambiguous structures, which may be partially or completely unstructured in solution, but can also assume a more-defined ordered structure in its specific functional state (structural folding on binding as observed in typical IDPs). Consistent with this, in native Raman and FT-IR spectrum of ExoY, abundant semi-ordered and molten conformers were also observed (Fig 2 **& S2, Table S1 & S2**). The lower abundance of β-structures, the propensity of T3SS effectors for initiation of amyloid-like aggregates during the unfolding of effector proteins seems to be reduced (52). In addition, the positively evolved proline residues may also contribute to effector unfolding, stability, and prevention of amyloid-like aggregation (53) (**Fig. S13**).

Another end of the unique compositional chemistry is the reduced overall residue-residue interaction potential (−7.76 v/s −12.61 of control bacterial proteome) in the sequence landscape of T3SS effector proteins (Fig. 6G-I, **Supporting Data 1**). This is essentially due to significantly decreased frequencies of hydrophobic or non-polar amino acids which provide major stabilizing interactions, and increased frequency of polar amino acids (such as serine and proline) which have significantly higher unfavorable interaction potentials (Fig. 6A-B, **Fig. S18, Supporting Data 1**). From residue-residue interaction potentials analysis, it is also clear that arginine residues can form more favorable stabilizing interactions than lysine residues (**Fig. S18**). Given that the frequency of arginine did not increase significantly compared to other polar or charged amino acids, whereas the frequency of another positively charged amino acid i.e. lysine increased more significantly in T3SS effector (possibly to maintain balance in charge distribution) (Fig. 6A & B). It seems clear that T3SS effectors are under strong evolutionary selection to minimize overall residue-residue contacts, folding and stability. Consistently, it can also be seen that the frequency of cysteine which forms a disulfide bond, one of strongest stabilizing interactions in proteins that require special enzymes or conditions to break, is significantly lower in T3SS effectors (Fig. 6B). Apart from serving a critical role in protein folding and stability, as non-specific hydrophobic collapse or non-polar interactions is also an eminent factor responsible for faster folding rates (34, 54). Underrepresentation of non-polar amino acids may plausibly contribute to slower folding rates in T3SS effectors (Fig. 6A-B). In addition, as geometrical frustrations arising electrostatic interactions are well known to reduce the number of favorable folding paths and folding rates (54). The increased unfavorable or repulsive polar-polar interactions propensities in polypeptide sequence landscapes will also contribute to slower folding kinetic in T3SS effectors (Fig. 6G-I).

Like arginine, even though histidine can form more favorable interactions (**Fig. S18**), interestingly, the frequency of histidine is significantly higher in T3SS effectors (Fig. 6A & B). Given that protonation of histidine (H46) played a critical role in mediating pH-dependent unfolding of T3SS effector AvrPto(8). Similarly, in ExoY, the histidine residues (H104, H46, H290, and H291) are among few polar interacting residues which play a critical role in maintaining labile tertiary structural stability (Fig. 7, **Fig. S11 & S12**). The positive evolution of histidine in T3SS effectors (Fig. 6A & B), despite its stabilizing nature **(Fig S18**), clearly suggests its adaptive role in PMF-mediated effector unfolding and secretion. To further affirm such imperative roles, in-silico mutational analysis of 104 histidine residues from 16 T3SS effectors was carried out (**Fig. S17, Supporting Data 1**). From our analysis, it could be seen that nearly 44.59 percent of histidine mutations are destabilizing, whereas the rest of 55.41 percent of mutations had neutral or stabilizing effects. At lower pH of 5.8, the destabilizing mutations significantly decrease from 44.6 to 36.9 %, while stabilizing and neutral mutations increased from 26.3 to 30.8 % and from 29.1 to 32.3 %, respectively (Fig. 7, **Fig S17)**. Thus, suggesting that the histidine residues not only play a vital role in folding and stability of T3SS effectors but can also tweak structural intricacy of effectors in presence of mild acidic conditions (based on increased stabilizing mutations of protonated histidine v/s non-protonated histidine residues), facilitating effector unfolding and secretion.

**Figure 7.**
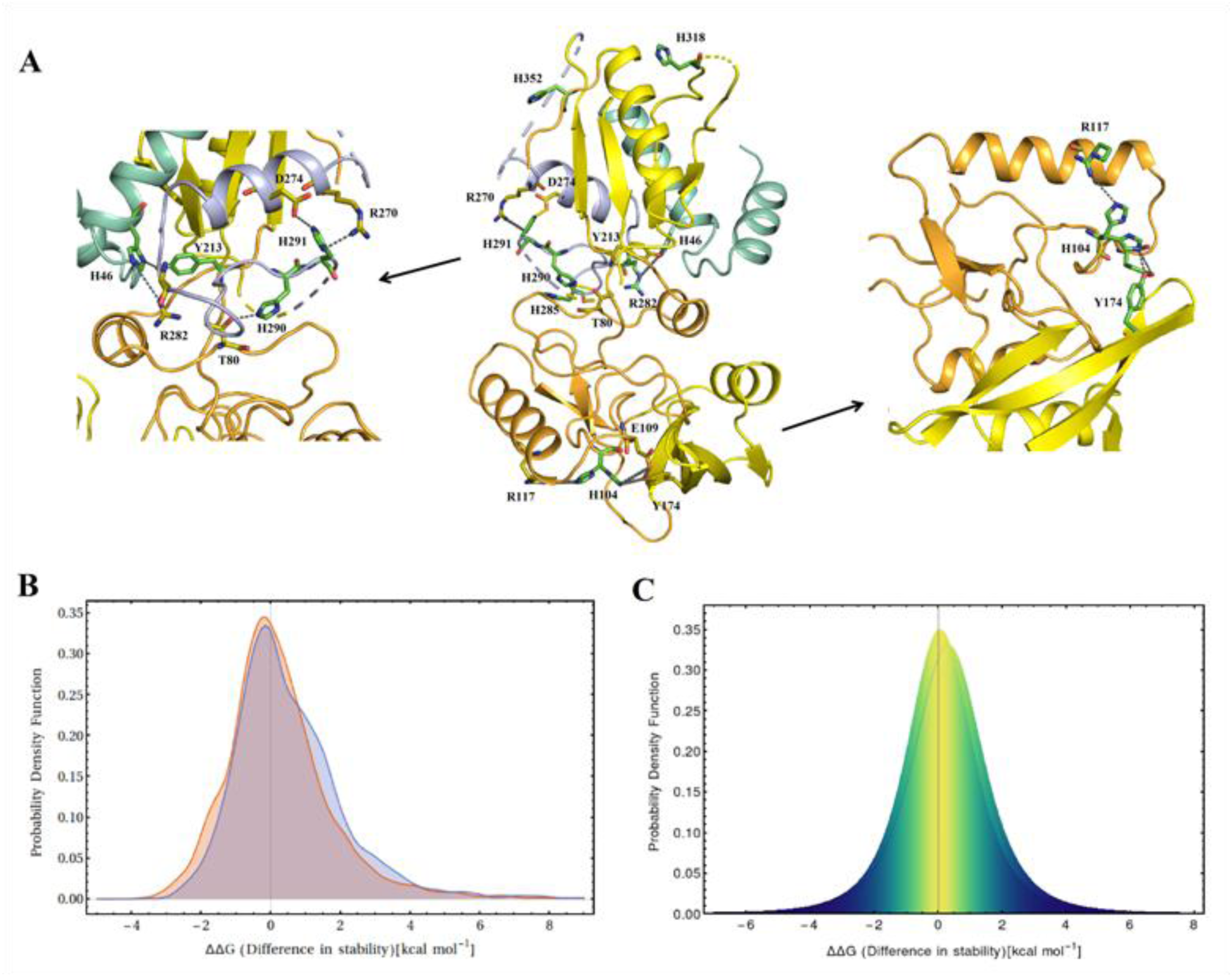
Site-specific evolutionary conservation of histidine residues drives a differential effect on the stability of T3SS effectors at physiological pH 7.4 and a mildly acidic condition of pH 5.8. **(A)** The overall distribution of histidine residues in the three-dimensional structure of ExoY (middle). The histidine residue (H46) of helix α2 formed salt bridge interaction with terminal tyrosine residue (Y213) of the barrel and terminal arginine residue (R282) of loop preceding helix α7. Similarly, the histidine residues (H290 and H291) from preceding loop of helix α7 had interacted with residue (T80) of disorder globule of ExoY and with residues (R270 and D274) of helix α7 and thereby binding central disordered globule with folded sub-domain 2. In sub-domain 1, histidine (H104) in disordered globule formed H-bonding interaction arginine residue (R117) of the helix (α5). Simultaneously with aid of glutamate residue from disordered globule, it also formed H-bonding with tyrosine residues (Y174) of barrel structure. These histidine mediated interactions were crucial for linking isolated helices and loops in disorder globule with stably folded structures that essentially contributed to the overall structural stability of ExoY. Upon protonation at low pH (<pKa of imidazole moiety of histidine side chains) can lead to destabilization of protein structure especially in presence of positively charged neighboring residues. The histidine residues in T3SS effectors can act as potential sensor cum switch for sensing low pH-mediated by proton-motive force (PMF) that can subsequently drive destabilization/unfolding of effector facilitating its secretion. To generalize such role of histidine residues in T3SS secretion, stability effects of mutation of histidine residues in T3SS effectors were investigated. The probability distribution of stability effect of histidine mutation was shown in figure **(B) (**obtained by Student T Distribution) and in figure **(C)** (obtained by kernel smoothing). The shift in the center of the probability distribution from more destabilizing effects at pH 7.4 to neutral and lesser destabilizing effects at pH 5.8 highlighted the more potential destabilizing nature of histidine residues at low pH than at physiological pH. These observations clearly depicted the potential sensor role of histidine residues in the spatial-temporal regulation of folding and stability of T3SS effectors.

From a detailed analysis of various structural and folding parameters, it is clear that T3SS effectors exhibit limited overall structural stability. Furthermore, to assess global tendency of T3SS effector proteins to fold or remain partially or completely unfolded, distribution of T3SS effectors in protein folding phase diagram was analyzed. According to charged-hydrophobicity boundary conditions of Uversky plot (55) (mean net charge v/s net hydrophobicity), the most of T3SS effectors predisposed in a folded section of phase diagram (Fig. 6J). In case of Das-Pappu phase diagram (56) (fraction of positive charge v/s fraction of negative charge), the most of T3SS effectors lay in the boundary between globular proteins and strong polyampholytes IDPs (Fig. 6K). To gain better insight on the degree of foldness, without biases towards net charge or hydrophobicity alone, distribution of T3SS effectors in three-dimensional charged-hydrophobicity phase diagram (57) was analyzed. Interestingly, in a revised folding phase diagram, most of the T3SS effectors belong to the unique class of unfolded proteins which comprise disordered globules (Fig. 6L).

The propensity of T3SS effector proteins to form disordered globules mainly stems from their lower preference for strong hydrophobic residues and higher preference for polar LCR promoting residues (N, S, Q, and T) and order breaker proline residues (Fig 6A-D). In addition, in contrary to earlier predictions that the disordered globules are mainly formed due to the collapse of less hydrophobic polar tracts (50, 56, 57), in T3SS effectors, the hydrophobic residues also play a non-negligible role in disordered globules formation. This is mainly due to the enrichment of weak hydrophobic amino acids such as Ala and Leu (compared strong hydrophobic acids) and their equivalent distribution in disordered regions of effector proteins (Fig 6B & D). Furthermore, as effector proteins have not evolved oppositely charged residues to a greater extent compared to polar uncharged residues, and their distribution in ordered and disordered segments is nearly equivalent (Fig 6A-D). The tendency of effector proteins to form strong polyampholytes induced swollen coils geometry is relatively less (56, 57) (Fig. 6K). Thus, as a consequence of abundance of hydrophobic LCR promoting residues (Ala and Leu) and polar LCR promoting residues (N, S, Q, and T) which can trigger hydrophobic collapse and proline and G-P rich LCR promoting segments which restrict the local folding i.e. formation of ordered secondary structures (49, 51, 58), T3SS effectors show a propensity for disordered globule formations (Fig. 6L). An outcome of such unique structural chemistry and co-existence of irrevocable folding parameters i.e. hydrophobic residues and G-P rich LCR promoting residues in disordered globules, T3SS effectors encompass abundant geometrically frustrated non-native hydrophobic clusters, which is an ultimate fuel for the global unfolding of effector proteins.

## CONCLUSION

Many pathogenic bacteria use T3SS injectisome to inject virulence effectors or toxins directly into the cytosol of targeted host cells (3, 22, 24). Given that the protein unfolding requisite for secretion via nano-size pore of T3SS apparatus is an energetically unfavorable process, “How do pathogenic bacteria unfold and secrete hundreds of toxic proteins in seconds” remain largely unknown. From our in-depth analysis of pH-dependent folding and stability of ExoY, and computational analysis of folding, stability, and structural evolution in other T3SS effectors, it is clear that the characteristic structural features evolved by T3SS effectors also play an eminent role in effector unfolding and T3SS secretion. These structural archetypes include the weaker and/or less compact hydrophobic packing, an abundance of unordered or intermediately folded coil structures, few long-range tertiary contacts, histidine-mediated stabilizing interactions, increased backbone solvation and geometrical frustration in the folded cores of effector proteins.

The random collapse of polypeptide chain for the exclusion of hydrophobic side chains from the surrounding solvent is one of the key events in protein folding (34), and solvating the buried hydrophobic core or structures is also a major barrier for the unfolding of proteins (32, 33, 36). From the absence of compact hydrophobic packing with the center of masses and lower abundance of non-polar or hydrophobic residues in effector proteome, it can be curtained that the hydrophobic packing in T3SS effectors is relatively weak and less compact. In addition, as a domain which encompasses more ordered secondary structural elements in ExoY shows more dispersed hydrophobic residues or clusters in the toroidal shape whereas domain comprising disordered globule with few ordered secondary structures show more compact hydrophobic packing. The co-evolution of compact hydrophobic packing and abundant ordered secondary structures that can confer higher tertiary structural stability seems to be avoided clearly in T3SS effectors. Furthermore, non-specific hydrophobic collapse or non-polar interactions, apart from serving a critical role in protein folding and stability, is also an eminent factor responsible for faster folding rates (34, 54). Underrepresentation of non-polar amino acids plausibly contributes to slower folding rates in T3SS effectors, an important trait essential for T3SS secretion. In addition, as geometrical frustrations arising from electrostatic interactions are well known to reduce the number of favorable folding paths and folding rates (54). The increased unfavorable or repulsive polar-polar interaction propensities in polypeptide sequence landscapes will also contribute to slower folding kinetic in T3SS effectors.

Consistent with earlier studies on Phytopathogenic effectors (59, 60), intrinsic disorderness is another conserved structural archetypes in all T3SS effectors and it mostly stems from abundance of highly polar and LCR promoting residues (such as N, S, Q, and T), charged residues (such as D, E, and K) and order breaker proline residues in their sequence landscapes (49, 50, 61). More, from our in-depth analysis, it can be also curtained that T3SS effectors belong to the typical class of IDPs which comprise disordered globules. However, in contrary to earlier predictions that the disordered globules are mainly formed due to the collapse of less hydrophobic polar tracts (50, 56, 57), in T3SS effectors, the hydrophobic residues also play a non-negligible role in disordered globules formation. This is mainly due to the enrichment of weak hydrophobic amino acids such as Ala and Leu (compared strong hydrophobic acids) in sequence landscapes of T3SS effectors proteins and their equivalent distribution in ordered and disordered regions. Furthermore, as a consequence of irrevocable folding parameters in the disordered globules, the T3SS effector proteins will encompass abundant non-native hydrophobic clusters and geometrical frustrations which ultimately act as a fuel for effector protein unfolding. Furthermore, as a consequence of unique structural chemistry and labile tertiary structural folding, slightest perturbation in physiochemical conditions such as a change in H^+^ or Na^+^ concentration can significantly elevate geometrical frustrations in effector proteins that eventually leads to partial unfolding of effector proteins. Consistent with this, T3SS export apparatus can unfold and secrete effector proteins in absence of ATPase and PMF, using Na+ concentration gradient (62). Moreover, the PMF-mediated unfolding of ExoY was energetically less unfavorable mainly due to the existence of solvated non-native hydrophobic clusters and abundance of geometrical frustrations in the folded cores.

Given that the secreted effectors proteins are very few when compared to their counterpart host proteins, the stability of effectors is also vital for their ability to function inside the host cell, which in turn determines the evolutionary fate of virulence effectors and pathogenic bacteria. One of the finest strategies for appeasing the contrasting needs is spatial-temporal regulation of effector stability to maintain balance in protein stability and function trade-off. PMF-mediated unfolding or destabilization of effector proteins is one of the finest strategies evolved by pathogenic bacteria to maintain balance in protein stability and function trade-off. From the evidences that the histidine residue, despite its stabilizing effect, is positively evolved in T3SS effectors, and plays an eminent role in folding and stability of AvrPto (8), ExoY, and other T3SS effectors, it is clear that histidine-mediated stabilizing interactions in T3SS effectors plausibly act as pH-sensing folding switches to regulate effector protein folding and stability in a spatial-temporal manner. In addition, T3SS effector proteins have evolved abundant gate-keeper residues and LCR promoting residues in their sequence space to prevent initiation of amyloid-like aggregation and to maintain a fine balance in folding, solubility, and stability (48, 49, 53). Lastly, from a detailed analysis of all facts and phenomena, it can be curtained that T3SS effectors belong to typical class (disorder globules) of IDPs and have evolved intrinsic disordered properties to facilitate their unfolding and secretion via nano-size pore of T3SS injectisome.

## MATERIALS AND METHODS

### Sample Preparations

ExoY was cloned, expressed and purified according to the previously reported methodologies(38). The expression of N^15^ and C^13^ labeled ExoY was carried by growing *E. coli* BL21 pLys harboring ExoY-pet28+ plasmid in a 5 mL LB media with appropriate antibiotic for 10 hours at 37 °C. The cells were then inoculated into 50 mL of M9 media containing radiolabel NH_4_Cl and Glucose, for overnight growth. The resulting cell culture was gently pelleted, resuspended in 1000 ml M9 minimal media prepared as above and grown till OD at 600 nm reaches 0.8-1.0. Then, the culture was transferred to 18 °C shaker, and protein over-expression was induced with 0.5 mM isopropyl-D-1-thiogalactopyranoside for 10 hours. The cells were harvested and the pellet was resuspended in lysis buffer containing 50 mM Phosphate Buffer pH 8, 150 mM NaCl, 10% Glycerol, 0.5 mM TCEP and 1X protease inhibitor cocktail. The cells were then lysed by sonication and protein was purified by Ni-nitrilotriacetic acid (NTA) chromatography. The eluted proteins from Ni-NTA column were loaded onto a Hi-load 16/60 Superdex 200 (GE Healthcare) gel filtration column pre-equilibrated with 25mM Tris pH 8.0, 150 mM NaCl, and 0.5mM TCEP. SEC was carried out using the AKTA prime Plus purification system with a flow rate of 1 ml/min and the collected fractions were analyzed by SDS-PAGE analysis.

### CD Spectroscopy

All the Circular dichroism (CD) measurements were performed using JASCO J-815 CD spectrometer equipped with a Peltier temperature control unit. The secondary structural signature spectrum in the region between 190 to 250 nm for each protein sample (2.4 µM) was recorded in a quartz cuvette with a path length of 1 mm. The CD parameters such data pitch, bandwidth, and scanning speed were set to 1 nm, 1 nm, and 20 nm/minutes, respectively. A total of 3-4 different scans (within 600 HT voltage range) for each protein sample was collected and subsequently averaged to get the final spectrum. The melting temperature of 20 µM ExoY at various pH conditions was estimated by measuring the change in CD ellipticity at 222 nm increasing temperature (10-90°C). The fraction of folded (*pF*) or unfolded (*pU*) population at a given temperature data point in melting curves was determined using a standard fitting analysis as described below.

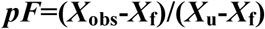

Where *X*_obs_ is observed CD ellipticity signal (mdeg) for each temperature data points, and *X*_u_ and *X*_f_ are CD ellipticity signal (mdeg) for fully unfolded and folded protein sample, extrapolated from linear fits of the melting curve regions.

### Raman Crystallography

The crystallization of ExoY was carried out according to the previously reported methodologies(38). In brief, the purified ExoY from gel filtration was combined with Elastase (0.1 mg/ml of protease for every 50 mg of protein) and then incubated at 15°C for 24-36 hours. After incubation, small degradation fragments were removed by concentrating ExoY using Amicon Ultra centrifugal filters (10 kDa cutoff). Finally, crystallization of ExoY was conducted in a sitting-drop vapor-diffusion system at 22°C by mixing 2 μl of protein solution (15-18 mg/ml) with 2 μl of reservoirs buffer containing 0.1M HEPES pH 7.5 and 1.4 M LiSO4. Well-defined tetragonal shaped proteins crystals were obtained after 15 - 20 days. Raman spectroscopic analysis of ExoY crystals was conducted using an STR Raman system (AIRIX Corp, Japan). The harvested protein crystals of ExoY were transferred to glass cover slip containing reservoir solution with pH of 5.8 and were incubated for 30-60 minutes. Finally, the incubated protein crystals obtained at 30 minutes and 60 minutes of intervals of incubation period were shot with 18.5 mW radiant power He-Ne laser (LASOS-LGK7665P18) obtained after excitation at 633 nm and spectra were recorded from 800-4000 cm^−1^ in spectral range with a data acquisition time of 150 sec. GRAMS/A1 software was used for processing Raman spectra.

### NMR Spectroscopy

NMR experiments were carried out on uniformly ^13^C & ^15^N doubled-labeled ExoY protein at 15°C in 20 mM sodium phosphate buffer (pH 7.4) and 10% D_2_O was used for locking purpose. All the NMR spectra including 2D ^1^H-^15^N HSQC and 3D triple resonance HNCA was conducted at 15°C on AV700 Bruker (^1^H resonance frequency 700 MHz) using a cryogenically cooled probe. At such high concentration (300 µM) inside NMR tube, ExoY suffered from gradual unfolding and phase separation during the time of recording of others 3D-triple resonance experiments. So, we could only perform HNCA data reliably and obtained reporting partial assignment unambiguously. ^1^H Chemical shifts calibration was executed using standard 2,2-dimethyl-2-silapentane-5-sulfonate (DSS) whereas the calibration of the resonance frequency of ^13^C & ^15^N chemical shifts were done indirectly. 2D ^1^H-^15^N HSQC experiment was conducted with (256 × 2048) complex points along the ^15^N and ^1^H dimensions, with an NMR data acquisition time of 90ms and 146ms respectively. Spectral windows and centered positions for ^1^H nucleus and ^15^N nucleus were 7002 Hz & 4.681 ppm and 1419 Hz & 118 ppm for ^1^H-^15^N HSQC experiments, respectively. 3D HNCA experiment was performed with a (64 × 128) complex points along the ^15^N and ^13^C dimensions and 2048 real data points in ^1^H dimension, with spectral widths and acquisition times for ^1^H, ^15^N and ^13^C nucleus were (9765.6 Hz & 104.9 ms),(1419 Hz & 22.55 ms) and (4929.6 Hz & 12.98 ms) respectively. All the NMR data processing was performed with TOPSPIN 4.0.3 software and backbone assignments were analyzed using CARA.

### Fourier Transform Infrared (FT-IR) spectroscopy

Attenuated total reflection infrared (ATR-FTIR) spectra of ExoY at different pH conditions were recorded using Bruker Tensor 27 spectrometer. Prior to the recording of FT-IR spectra, protein samples were incubated at various pH conditions for at least 2 hours. The respective buffer baseline was subtracted according to previously established double subtraction procedures(31). The assignment of spectral components was done according to the earlier reports describing spectral components associated with different secondary structure(29, 30). For deuterium/hydrogen amide exchange kinetics, 50 µl of 186 µm of ExoY was mixed with 200 µl of deuterated buffer solutions. All the buffers were prepared in D_2_O. For every spectrum, at least 60-scans interferograms in a single-beam mode with 4 cm^−1^ resolution were recorded for almost 40 minutes after every 150 seconds. However, for analysis, due to intrinsic instability of ExoY at room temperature spectra beyond 18 minutes were excluded. Similarly, reference spectra were also recorded under identical scan conditions. Finally, each of the spectra was processed according to previously established double subtraction procedures.

### Fluorescence spectroscopy

All the fluorescence spectra were collected on Carey Eclipse Fluorimeter (Sl.no.MY13130004) using 1ml quartz cuvette (Hellma Analytics) with a path length of 10 mm. Scan speed was configured to 100 nm/min using response time of 1 sec and slit width of 4-7 nm for excitation and emission. The protein concentration used for all fluorescence spectra accumulation was 5 µM otherwise mentioned whenever necessary. All the spectra shown were obtained after subtracting with a proper blank. Intrinsic tryptophan fluorescence of ExoY to monitor local unfolding was carried out at various pH conditions by selectively exciting tryptophan at 295 and recording emission spectra between 305 to 450 nm. Similarly, the *bis-*ANS fluorescent dye binding assay, to monitor the exposure of hydrophobic surfaces was carried out by selectively at exciting *bis*-ANS at 385 nm and collecting the emission spectrum between 400 to 600 nm. Around 20 µl of 1 mM *bis*-ANS was for every 1ml of 5 µM of protein samples. To monitor the planner β-pleated sheet conformation in coacervated species or self-assembled states of ExoY, ThT binding assay was performed. Around 20 µl of 2 mM ThT dye for was mixed properly with 1 ml of 5 µM protein samples. ThT in buffer without protein was used as a baseline. ThT was specifically excited at 440 nm and emission spectra were acquired from 460 to 600 nm with integration time was of 1 sec.

### Isothermal Titration Calorimetry

To determine the energetics of the PMF-mediated acid unfolding of ExoY, Isothermal titration calorimetric (ITC) experiments were performed using a Microcal VP-ITC titration calorimeter (MicroCal, LLC, and Northampton, MA). A sample cell containing 1 ml of 17 µM ExoY in 1mM Phosphate buffer pH 7.8, 50 mM NaCl, and 0.01 mM TCEP was titrated with 4 mM HCl (syringe) with the constant stirring speed of 25 rpm’s at a temperature of 15°C. A total of 27, 1 µL injections were made every 180 seconds to decreases the pH from 7.8 to 5.6. All the samples were extensively degassed before performing titration experiments. The resulting titration curves were processed using the Origin ITC software package supplied by MicroCal (Northampton, MA) to obtain a change in enthalpy of reaction (Δ*H*°) and change in entropy of reaction (Δ*S*°). Finally, the standard Gibbs free energy change (Δ*G°*) was calculated using the following equation: (Δ*G°* = Δ*H*°-TΔ*S*°)

### Microscopic Imaging

Phase separation and formation of coacervated species of full-length and truncated ExoY (after proteolysis) was monitored by using an inverted optical microscope (Laika DMI 4000, Leica and Zeiss Co., Cambridge, England) equipped with a motorized stage. Phase separation of ExoY was triggered by incubating at room temperature (25°C) for 5-10 minutes. Around 5-10 µL of protein sample (10-20 µM) was drop cast on clean glass-slide and was covered with a clean coverslip. Optical images were then recorded through QWin software using illumination from the mercury lamp or the laser. Transmission Electron Microscopy (TEM) analysis of coacervated species was performed by a JEOL 1200EX transmission EM operating with an accelerating voltage of 120 kV. 3-5 µL of a stock solution of protein sample (0.5 µM) was drop cast on 400 mesh copper grids (PST product, GSCu300C-50). After 3-5 minutes, an excess of the sample was adsorbed to the dust-free filter paper. Grids were gently rinsed with MilliQ water and finally stained with 1% Uranyl acetate for 25 min at room temperature.

### Bioinformatics & Computational Analysis

For statistical analysis of conserved physiochemical properties in T3SS effectors, a local database of 371 confirmed T3SS effectors from 18 bacterial genus and 25 bacterial species was generated from the text-based literature search. Further, to avoid any phylogenetic biases, the conserved secreted effectors such early secreted translocator and needle tip proteins were filtered. Lastly, the protein sequences of each T3SS effector from a final list of 275 T3SS mediated secreted effectors, were retrieved from UniProt/NCBI database. For amino acid composition analysis, 254 effectors with no sequence and structural homologies among themselves were used. The amino acid frequency in T3SS harboring bacterial proteomes was obtained from previously published datasets(63) and was subsequently used as a control for comparative analysis.

The pairwise residue-residue interaction potential energies were estimated using the following equation(61, 64):

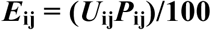

Where, *U*_ij_ is the inter-residue contact potential energy between i and j residue types, and *P*_ij_ *is the* probability of their interaction in the protein sequence, which is based on the frequency of occurrence of i and j residue. Inter-residues contact potential energies were obtained from the previous studies(65). Further, based on the additivity of the energy, the overall residue-residue interaction energies for T3SS effector and control proteome, and the difference in overall residue-residue interaction energies for T3SS effector and control proteome were also calculated.

The instability index, aliphatic index and gravy index of effectors were computed using ExPASy-Protpram tool. The secondary structure composition in T3SS effectors was computed using SPIDER2 algorithm(66). Similarly, the disordered residues in effector proteins were estimated using GeneSilico MetaDisorder server(67). Further, secondary structure and disordered predication were superimposed and overall distribution of helices, strands, coils and disordered residues in effectors was estimated by counting the number of residues encoding helix, strand, coil, and disordered residues. The distribution of T3SS effectors in the diagram of protein folding states proposed by Uversky plot(55) and Das-Pappu phase(56) was carried using localcider package(68). The role of particular amino acid on folding and thermodynamic stability of protein was probed using in-silico mutational analysis tool of FoldX(69) (version 2.52). The analysis comprised of four simple steps as described in detail previously(69, 70). In brief, first, the three-dimensional crystal structure of protein retrieved from the Protein Data Bank was optimized using the repair function of FoldX. Secondly, using the position scan repair function of FoldX, multiple structures of single point mutants (including self-mutated structures) were created. Third, using the energy calculation function of FoldX, the difference in free energy of wild-type and mutant structures was computed at two different pHs i.e. pH 7.4 and pH 5.8. Lastly, the difference in free energy values of the mutant structures with those of the wild-type structures to obtain the ΔΔG values of mutations.

## FOTENOTES

Abbreviations used in manuscript: T3SS, Type-III secretion system; PMF, Proton motive force; NMR, Nuclear magnetic resonance; FT-IR, Fourier transform infrared spectroscopy; ITC, Isothermal titration calorimetry; CD, Circular dichroism; PPII, Poly-l-proline II; LCR, Low complexity regions; ANS, 8-Anilino-1-naphthalene sulfonic acid; ThT, Thioflavin T; IPTG, Isopropyl-β-D-thiogalactoside; NTA, Nitrilotriacetic acid

## ACKNOWLEDGMENTS

All the authors also thank Jishu Mandal, Abhishek Das, Chiranjit Biswas, Sandip Dolui and Bidisha Chakraborty for their assistance.

