## Supporting Information for "Deciphering the structural intricacy in virulence effectors for proton-motive force mediated unfolding and type-III protein secretion"

**Table S1.** Raman vibrational bands from 900-1800 $cm^{-1}$ of a single crystal of ExoY at pH 7.4 and 5.8

**Table S2.** FT-IR peaks assignment from 1500-1700 $cm^{-1}$ of ExoY at pH 7.4 and 5.8

**Table S3.** Distribution of ordered and unordered secondary structural elements in T3SS effectors

**Figure S1**. Sequence length distribution of T3SS effectors

**Figure S2.** Folding and stability of ExoY in presence of mild acidic conditions of proton-motive force

**Figure S3.** Raman crystallographic analyses of ExoY single crystals at pH 7.4 and 5.8

**Figure S4**. Fourier transform infrared (FT-IR) spectroscopic analysis of ExoY at pH 7.4 and 5.8

**Figure S5**. Envisaging tertiary structural unfolding of ExoY by monitoring intrinsic tryptophan fluorescence and global exposure of buried hydrophobic surfaces

**Figure S6.** Energetics of PMF-mediated acid unfolding of ExoY

**Figure S7.** Characterizing intrinsically disordered property of ExoY as a function of pH

**Figure S8.** Envisaging the pH-dependent global unfolding propensity of ExoY from the ability to form thermoresponsive coacervated species

**Figure S9.** 2D NMR spectra of ExoY

**Figure S10.** Distribution of disordered residue as detected from NMR experiments in the crystal structure of ExoY

**Figure S11.** Hydrophobic clustering in the crystal structure of ExoY

**Figure S12.** Hydrophobic packing and geometrical frustration in the folded cores of ExoY

**Figure S13.** Decoding the structural stability of folded sub-domain 2

**Figure S14.** Distribution of charged residues and long-range electrostatic interactions between oppositely charged residues in ExoY

**Figure S15**. Evolutionary impact of proline and glycine-proline enriched ELP-like peptide motifs in ExoY

**Figure S16**. Potentially optimized helix stabilizing and destabilizing interactions in isolated helices of ExoY

**Figure S17.** Geometrical stress in short amphipathic twisted β-bladed conformations embedded in the core of relatively stable barrel-like fold of ExoY

**Figure S18.** The hypothetical model for the mechanism of effector protein unfolding and T3SS secretion

**Figure S19.** Distribution of stability effects of in-silico mutagenesis of histidine residues in T3SS effectors

**Figure S20A.** Residue-residue interaction energies of charged amino acids (D, E, K, and R) in T3SS effectors and control bacterial proteome respectively

**Figure S20B.** Residue-residue interaction energies of polar amino acids (N, S, Q, and T) in T3SS effectors and control bacterial proteome respectively

**Figure S20C.** Residue-residue interaction energies of hydrophobic aromatic amino acids (F, Y, and W) and methionine (M) in T3SS effectors and control bacterial proteome respectively

**Figure S20D.** Residue-residue interaction energies of hydrophobic non-aromatic amino acids (A, V, L, & I) in T3SS effectors and control bacterial proteome respectively

**Figure S20E.** Residue-residue interaction energies of G, P, C, and H in T3SS effectors and control bacterial proteome respectively

**Supporting References**

**Table S1**. Raman vibrational bands from 900-1800 $cm^{-1}$ of single crystals of ExoY at pH 7.4 and 5.8

| pH 7.4 | pH 5.8 (30 min) | pH 5.8 (60 min) | Modes of Raman vibrations |
|---|---|---|---|
| 856 | 847 | 852 | Tyr |
| 904 | 910 | 886 | Cα–C stretching |
| 947 | 947 | 950 | helix skeletal, Cα–C stretching |
| 984 | 984 | - | P=O bending mode (symmetric) of phosphate ion |
| 1003 | 1003 | 1003 | Phe |
| 1096, 1112 | 1096, 1112 | 1112 | $\upsilon_{CC,}\upsilon_{CO,}$ $\upsilon_{CN}$ |
| 1124 | 1139 | 1145 | $\upsilon_{CC}$ |
| 1221, 1240 | 1215 | - | β-strand & sheet, extended PPII, amide III |
| - | 1251 | - | 3/10- helix, Amide III |
| 1258 | 1266 | 1265 | α-helix, amide III |
| 1322 | 1318 | 1326 | left handed PPII, amide III |
| 1345 | 1352 | 1350 | $C_{\alpha}$-H deformation, pure α-helix, amide III |
| 1449, 1460 | 1448, 1461 | 1458, 1470 | $CH_{2-}$,$CH_{3-}$& CH- deformation and scissoring |
| 1605 | 1614 | - | Phe, Tyr aromatic ring vibration |
| 1645 | 1648 | - | Turn/unordered, amide I |
| 1657 | - | 1654 | Helix, amide I |
| 1671 | 1670 | 1675 | Extended β-strand, amide I |
| - | 1681 | 1687 | Poly-l-proline II (PPII), random β-space, amide I |

**Table S2.** FT-IR peaks assignment from 1500-1700 $cm^{-1}$ of ExoY at pH 7.4 and 5.8

| $\nu_{cm}^{-1}$ (pH 7.4) | $\nu_{cm}^{-1}$ (pH 5.8) | Normal modes of vibrations |
|---|---|---|
| 1506, 1516 | 1506,1517 | $\nu_{C=C,}$ $\nu_{CH,}$ $\nu_{CH}$-bending and stretching |
| 1528,1536 | 1531 | $\nu_{C-N}$ stretching, $\nu_{N-H}$ bending, amide II |
| 1546,1550 | 1548,1554 | $\nu_{C=O}$ asymmetrical stretching, amide II |
| 1569,1582 | 1567,1582 | $\nu_{C=C}$ stretching, aromatic ring vibration |
| 1625 | 1626 | β-strand & sheet, amide I |
| 1633 | 1631 | Extended PPII-helix, β-strand, amide I |
| 1640 | 1639,1643 | (3/10)-helix, Poly-l-proline, amide I |
| 1648,1653 | 1648 | α-helix, amide I |
| 1662 | 1660,1676,1694 | β-strand/turns, amide I |

**Table S3.** Distribution of ordered and unordered secondary structural elements in T3SS effectors

| Bacterial Taxonomy (order) | No of effectors | SS-3C analysis | | | SS-4C analysis | | | |
|---|---|---|---|---|---|---|---|---|
| | | Coil (%) | Helix (%) | Sheets (%) | Disorder (%) | Coil (%) | Helix (%) | Sheets (%) |
| **Pseudomonads** | 78 | 49.67 | 39.01 | 11.32 | 42.75 | 21.72 | 26.69 | 8.84 |
| **Vibrionales** | 12 | 41.78 | 50.53 | 7.69 | 42.73 | 17.47 | 34.07 | 5.73 |
| **Chlymadiales** | 23 | 44.14 | 39.27 | 16.60 | 37.57 | 19.09 | 28.89 | 14.45 |
| **Xanthomonadales** | 45 | 48.24 | 44.67 | 7.10 | 45.65 | 18.52 | 30.96 | 4.88 |
| **Nisseriales** | 11 | 42.76 | 48.78 | 8.46 | 39.63 | 20.25 | 33.18 | 6.93 |
| **Rhizobiales** | 13 | 56.98 | 31.66 | 11.36 | 58.18 | 17.24 | 18.30 | 6.28 |
| **Aeromondales** | 9 | 40.11 | 50.36 | 9.53 | 50.51 | 15.29 | 27.54 | 6.66 |
| **Burkholderilaes** | 46 | 49.18 | 40.86 | 9.97 | 42.91 | 21.54 | 28.33 | 7.22 |
| **Enterobacteriales** | 134 | 46.76 | 41.52 | 11.73 | 29.65 | 28.09 | 32.19 | 10.08 |
| **Total** | 371 | 47.77 | 41.69 | 10.55 | 39.45 | 22.60 | 29.65 | 8.30 |

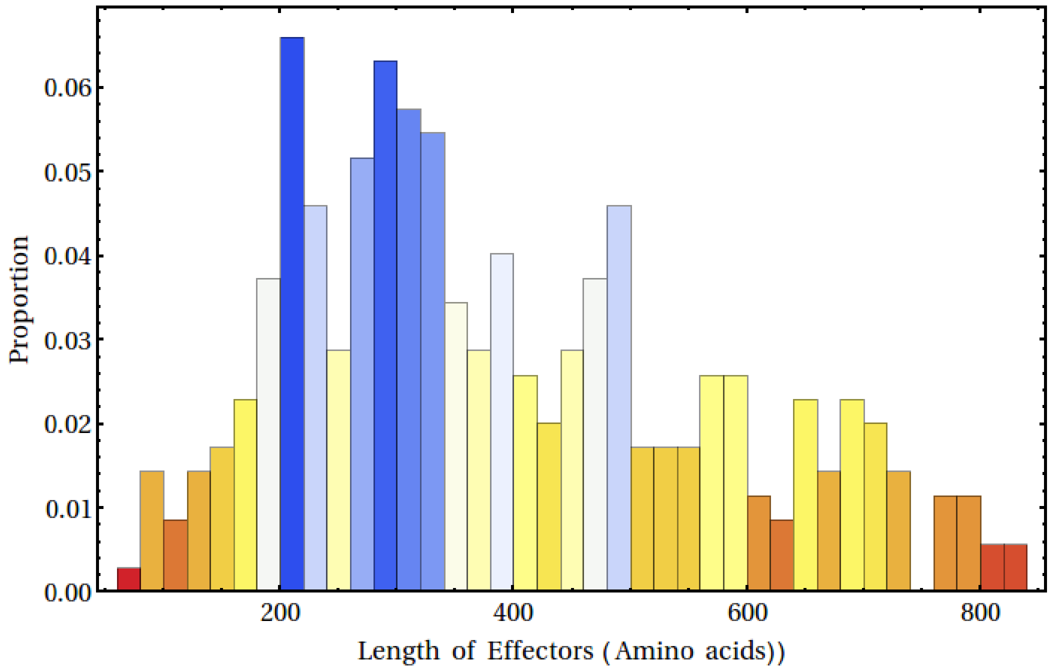


**Figure S1. Sequence length distribution of T3SS effectors.** The majority of T3SS effector proteins have more than 200 amino acids in their polypeptide sequence with mean size of 453 amino acids (~ 49 kDa)

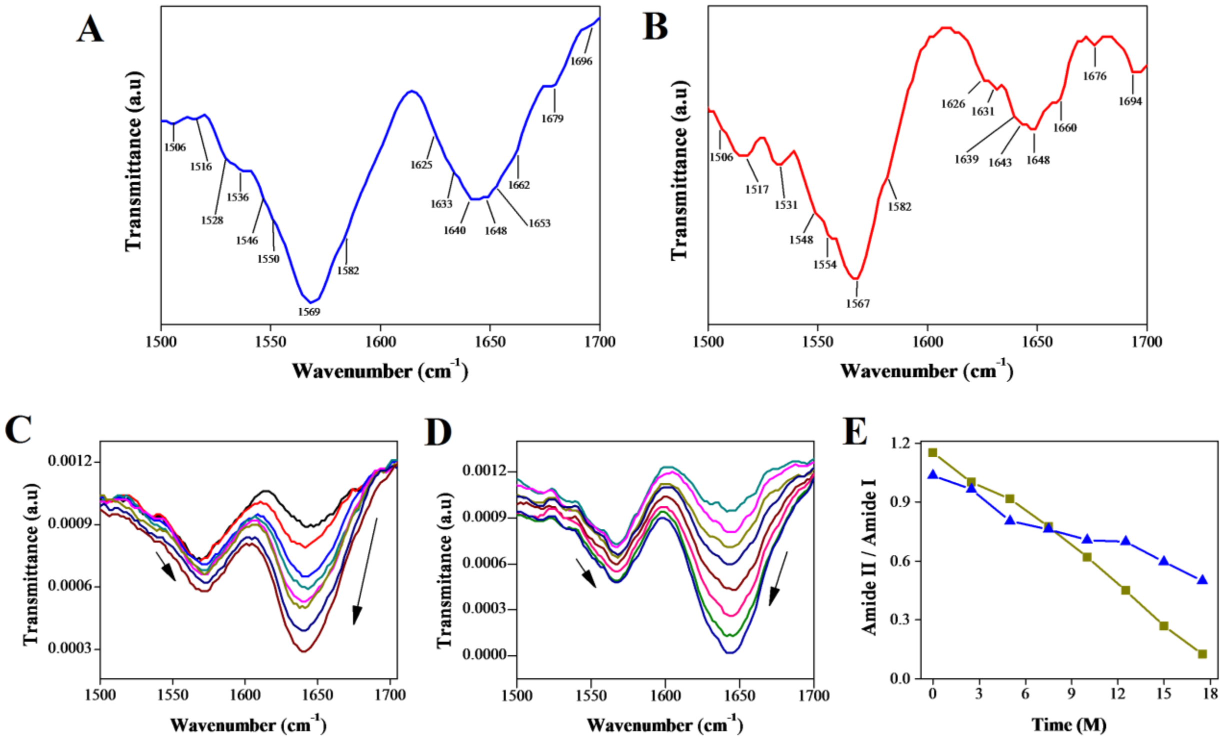


**Figure S2. Secondary structural characterization of ExoY at pH 7.4** and **5.8.** To understand spectral components associated with the different secondary structure, a partly assign FT-IR spectra from (1500-1700 cm$^{-1}$) containing amide I and amide II regions of full-length ExoY at **(A)** pH 7.4 **(B)** pH 5.8 was performed. FT-IR spectra (1500-1700 cm$^{-1}$) showing hydrogen-deuterium (H/D) exchange kinetic rates of ExoY at **(C)** pH 7.4, **(D)** pH 5.8. Arrows indicate amide I and amide II spectral changes as a function of time. **(E)** From time-course H/D exchange plot i.e. ratio of amide II to amide I (1550/1650) clearly show faster amide exchange rates in ExoY at pH 5.8 (red) compared to pH 7.4 (blue).

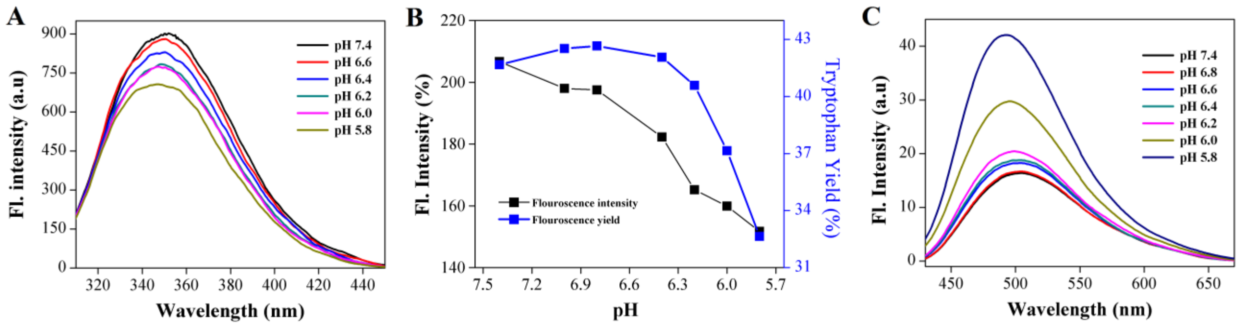


**Figure S3. Envisaging tertiary structural unfolding of ExoY by monitoring intrinsic tryptophan fluorescence and global exposure of buried hydrophobic surfaces. (A)** Gradual quenching of intrinsic tryptophan fluorescence indicates the exposure of buried tryptophan in ExoY with lowering of pH from 7.4 to 5.8. **(B)** Analysis of tryptophan fluorescence yield further accentuates the partial exposure of buried tryptophan to bulk solvent with critical transition era of pH 6.5-5.8 **(C)** Gradual and systematic increase in ANS fluorescence with a decrease in pH from pH 7.4 to 5.8 indicates a partial exposure of buried hydrophobic regions in ExoY.

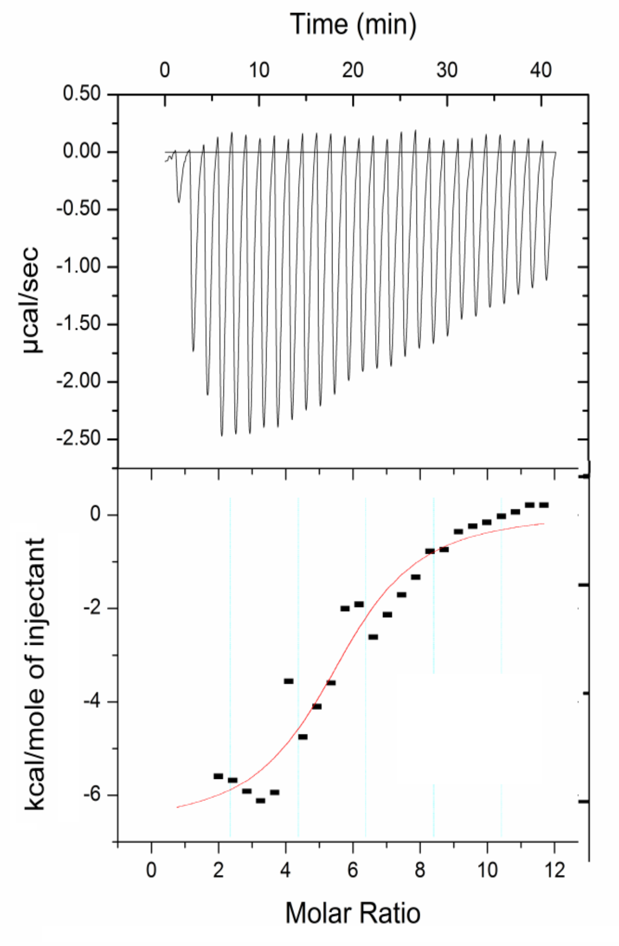


**Figure S4. Energetics of PMF-mediated unfolding of ExoY.** Isothermal calorimetric titration experiment indicate that the thermodynamic feasibility of PMF-mediated unfolding of ExoY PMF-mediated acid unfolding of ExoY is primarily enthalpically accelerated process ($\Delta H^{o}$ = - 6.4 Kcal/mol) with minimal entropic cost ($\Delta S^{o}$ = - 6.4 Kcal/mol). The standard Gibbs free energy ($\Delta G^{o}$ = -5.89 Kcal/mol) between native state (pH 7.8) and pH-induced partly unfolded state of ExoY (pH 5.6) also indicate the thermodynamic feasibility of PMF-mediated unfolding of ExoY. The highly favorable enthalpy factor with minimal entropic cost suggest the existence of partially solvent-exposed large polar and charged residues inside the labile tertiary structure of ExoY and PMF-mediated unfolding mainly involved exposure of these buried polar tracts. These observations are consistent with spectroscopic analyses which show dramatic alteration in tertiary structure and backbone solvation of ExoY, without significant increase in exposure of buried hydrophobic surfaces (**Fig 1 & 2, Fig S2 & S3**).

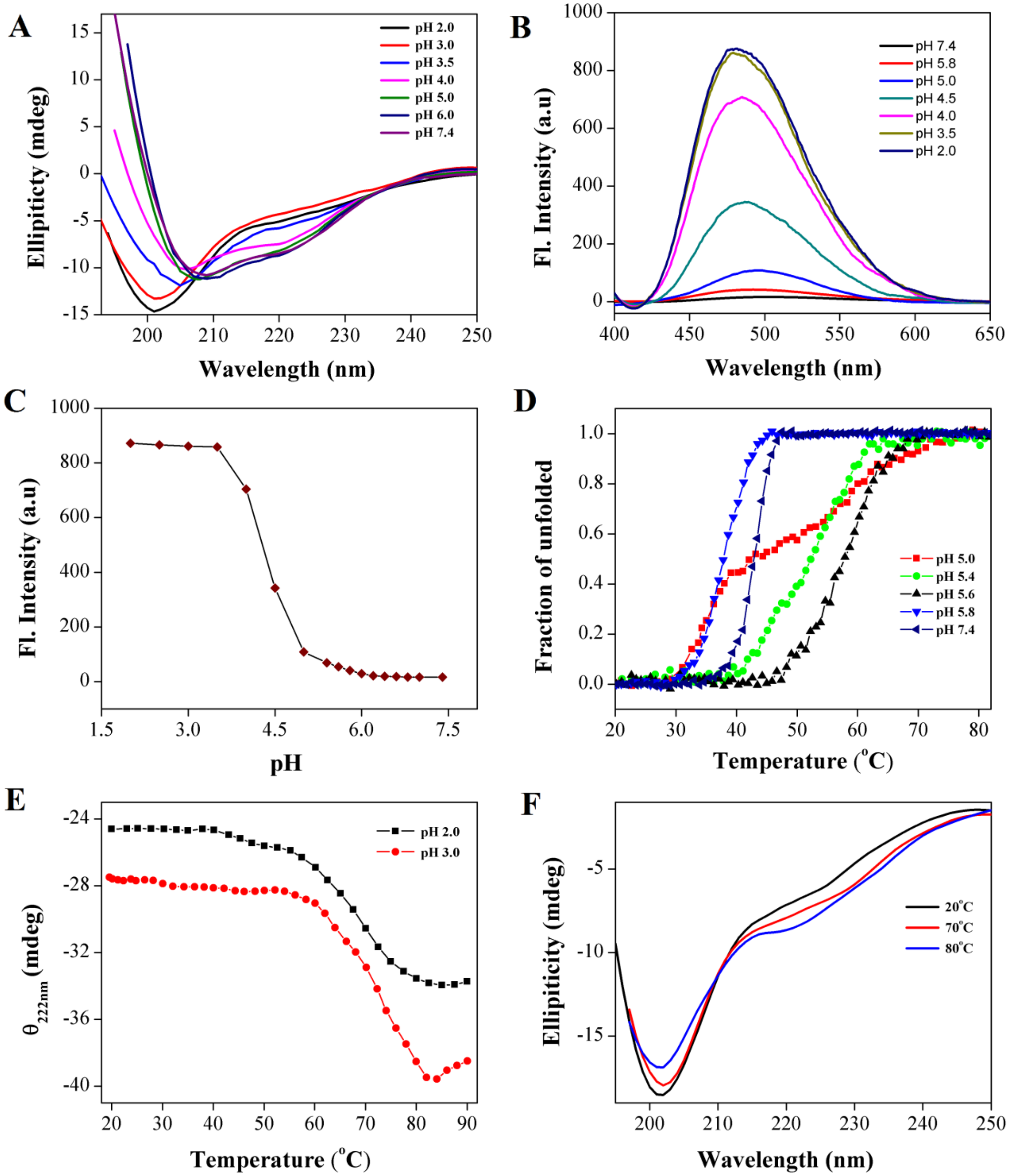


**Figure S5. Characterizing intrinsically disordered property of ExoY as a function of pH**. **(A)** Far UV-CD spectra showing gradual and complete secondary structural unfolding of ExoY at lower pH conditions. A dramatic loss in the secondary structure of ExoY could be primarily initiated only by lowering the pH below transition period of pH 5. **(B)** Monitoring the exposure of buried hydrophobic surfaces in ExoY during equilibrium acid unfolding experiments by ANS dye binding assays. **(C)** The observed ANS fluorescence at pH 5.8 is just 3.22 % of total fluorescence observed for completely acid unfolded ExoY. Similar to dramatic loss of folded secondary structural contents, exposure of buried hydrophobic surfaces in ExoY could be largely initiated after lowering the pH below 5 **(D)** Increased melting temperature or multiphase thermal melting curves of ExoY at pH 5.6-5.0 was due to partial exposure of buried hydrophobic cores and formation self-assembled non-native protein complexes. **(E)** At extremely lower pH 2 and 3, instead of equilibrium unfolding, ExoY shows temperature-induced structural compaction. **(F)** The increased ellipticity at 222 nm and decreased ellipticity at 208 nm in far-UV CD spectra of ExoY also indicates partial adaptation of helical conformer and loss of unordered secondary structures. Such unusual structural compaction or folding is a usual phenomenon of many intrinsically unstructured proteins due to inverted free-energy folding landscape(1–3).

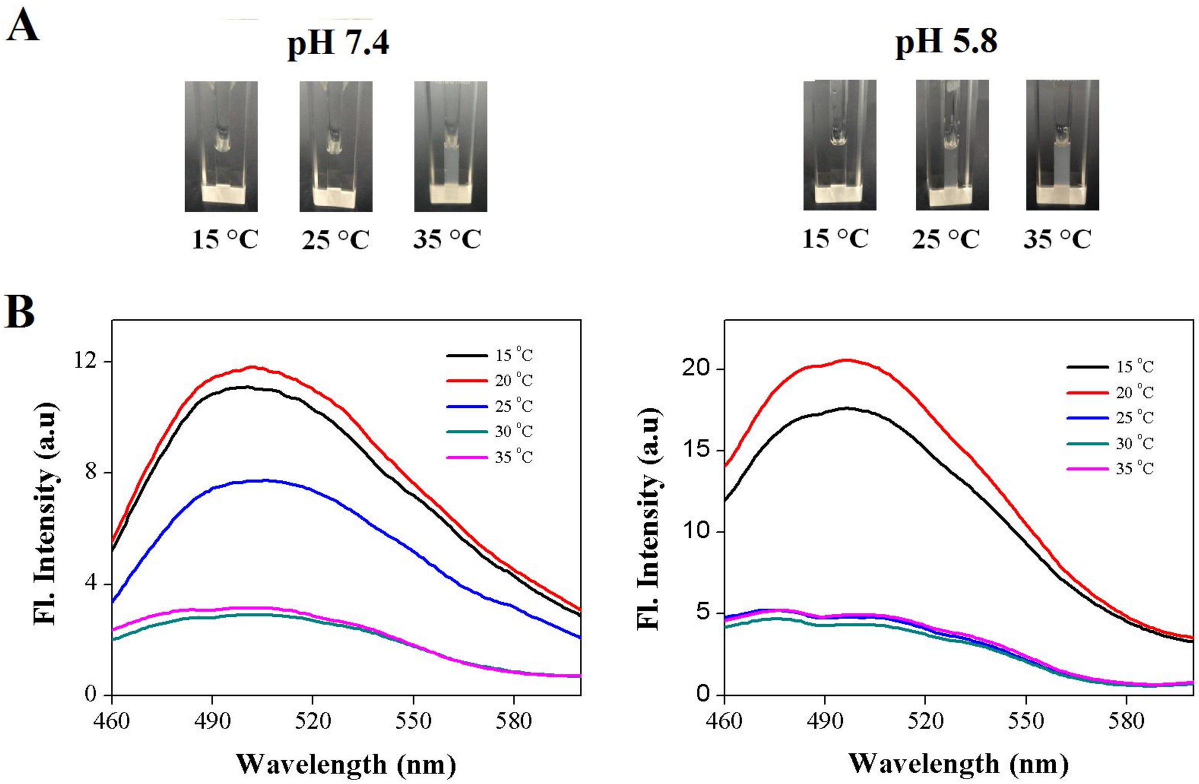


**Figure S6. Envisaging the pH-dependent global unfolding propensity of ExoY from the ability to form thermoresponsive coacervated species. (A)** Turbidity assay depicting the formation of condense liquid-liquid phase separated protein or coacervate of ExoY as a function of temperature and pH. Coacervation propensity of ExoY was relatively higher at pH 5.8 (right) compared to pH 7.4 (left). In presence of mild acidic conditions of PMF, the reduced critical temperature for formation of thermoresponsive coacervated species of ExoY depict the role of IDP-initiated tertiary structural unfolding in liquid-liquid phase separation(4, 5). **(B)** ThT-dye binding studies for detecting cross β-sheet structure formation in self-associated or coacervated species of ExoY. Partial loss in ThT fluorescence of ExoY at pH 7.4 (left) and pH 5.8 (right) (due to the scattering of insoluble condensed protein phase separated or coacervates species) after critical phase transition temperature (25$^{o}$C) suggested that the formation of thermoresponsive coacervates species is not due to the formation of typical cross β-sheet enriched amyloid-like aggregates(4, 6–8). ThT fluorescence scattering results also suggested the lowering of critical transition temperature at lower pH and the rapid formation of condense protein coacervates is mainly due to pH induced IDP-initiated tertiary structural unfolding or melting of disordered globule (rich in ELP-like peptide sequences) of ExoY(4, 5, 9).

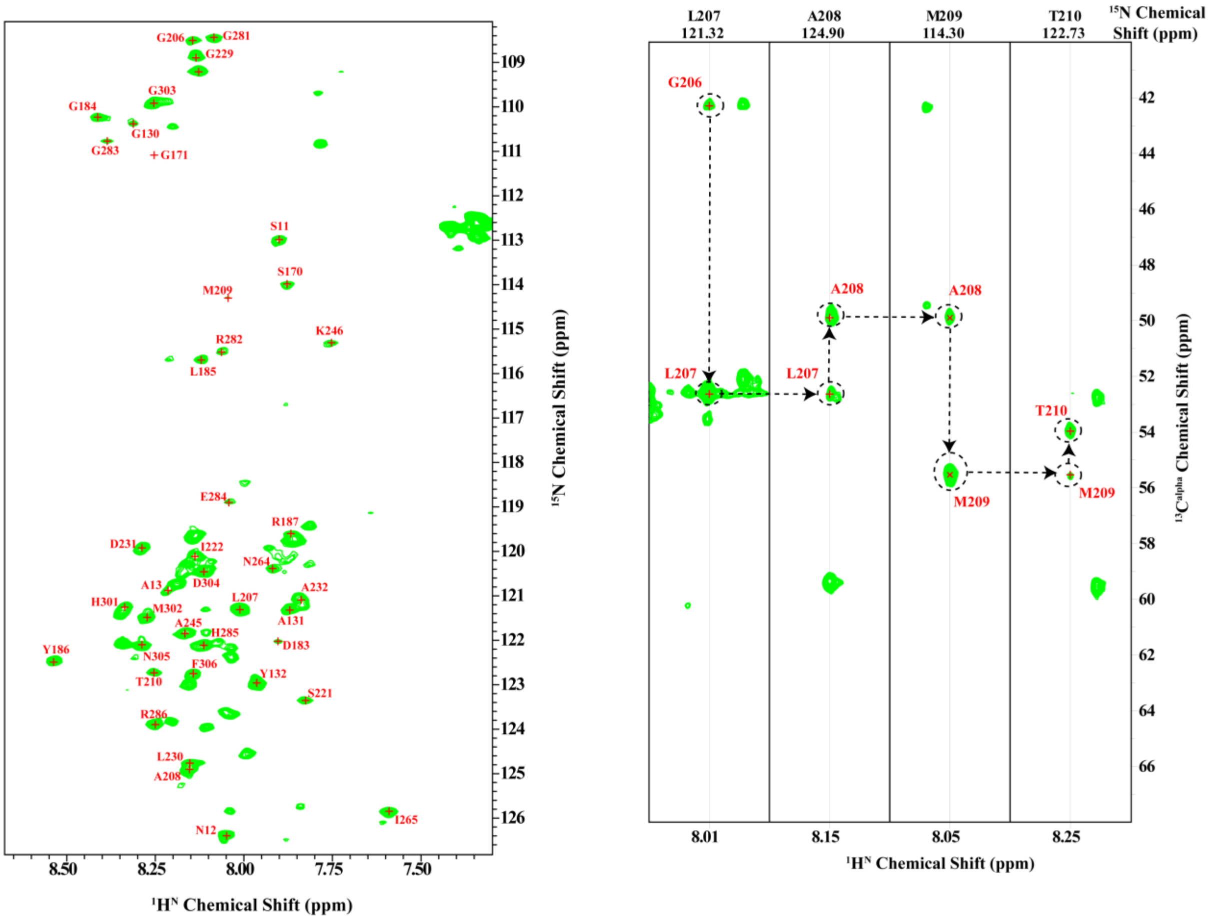


**Figure S7. 2D NMR spectra of effector protein ExoY (A)** Partially assigned 2D $^{1}$H-$^{15}$N heteronuclear single quantum spectrum (HSQC) of full-length ExoY (300 µM) at 15 $^{o}$C in 20 mM sodium phosphate buffer (pH 7.4). All the assigned peaks are shown in single letter code of amino acids. Due to the large deviation in the longitudinal relaxation time between folded and disordered amide protons, signals from mostly the disordered residues are primarily predominant in HSQC spectra as they have much sharper line-widths compared to folded counterparts. Except for large abundant proline and side chain residues from Asparagine and Glutamine, we observed 70 cross-resonance peaks from highly disordered residues. **(B)** Strip-plot in $^{15}$N plane of 3D HNCA spectra extracted from residues (G206 -T210) of ExoY protein and sequential spin connectivity was depicted by the dashed line as a representative case for HNCA based assignment strategy.

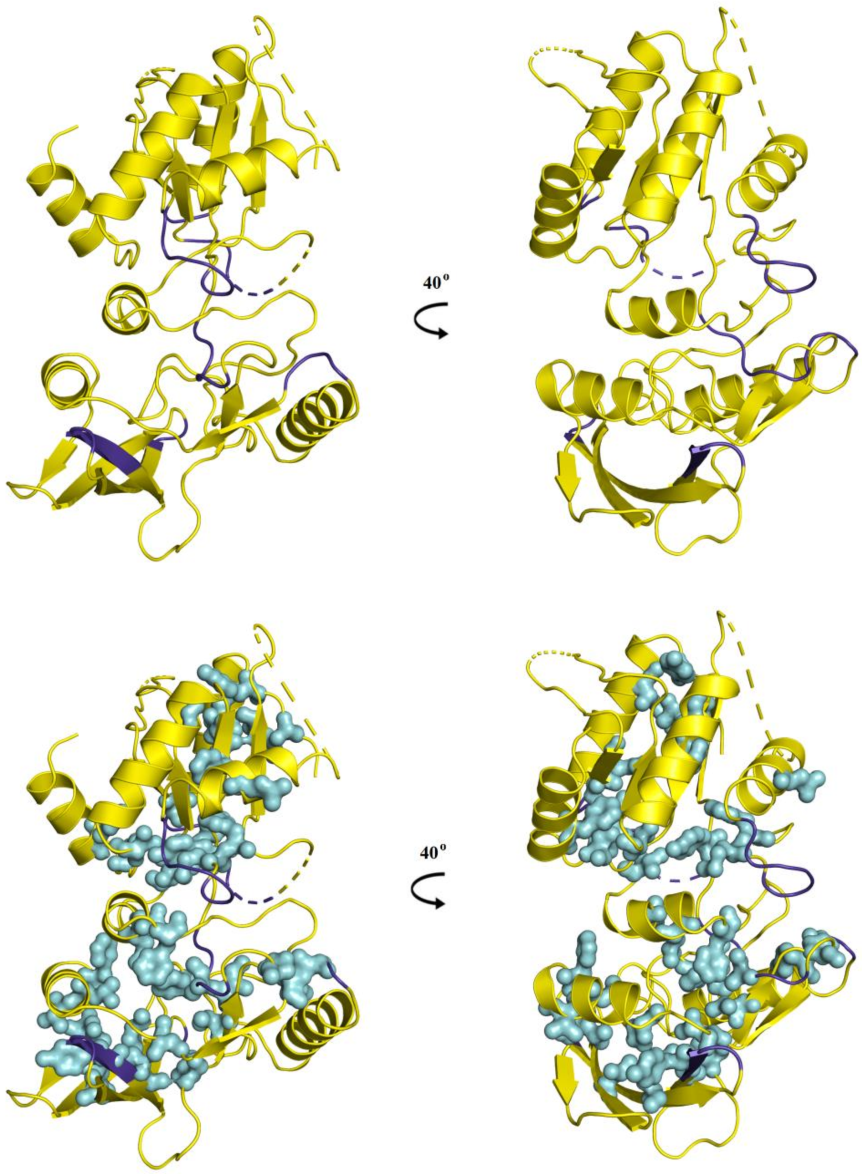


**Figure S8. Distribution of disordered residues as detected from NMR experiments in the crystal structure of ExoY.** Crystal structure of ExoY (PDB: 5XNW) showing disordered residues (marked in blue color) as observed in NMR experiments**.** Out of 42 assigned disordered residues, most of the assigned amino acids were found to distributed on surface exposed loops and a few were localized either in extended β-strand in the crystal structure or belonged to missing regions of the crystal structure. Interestingly, the NMR-detected disordered residues or regions were mostly surface exposed and did not co-exist with the residues that were localized in hydrophobic clusters of central disordered globule region (highlighted in cyan color).

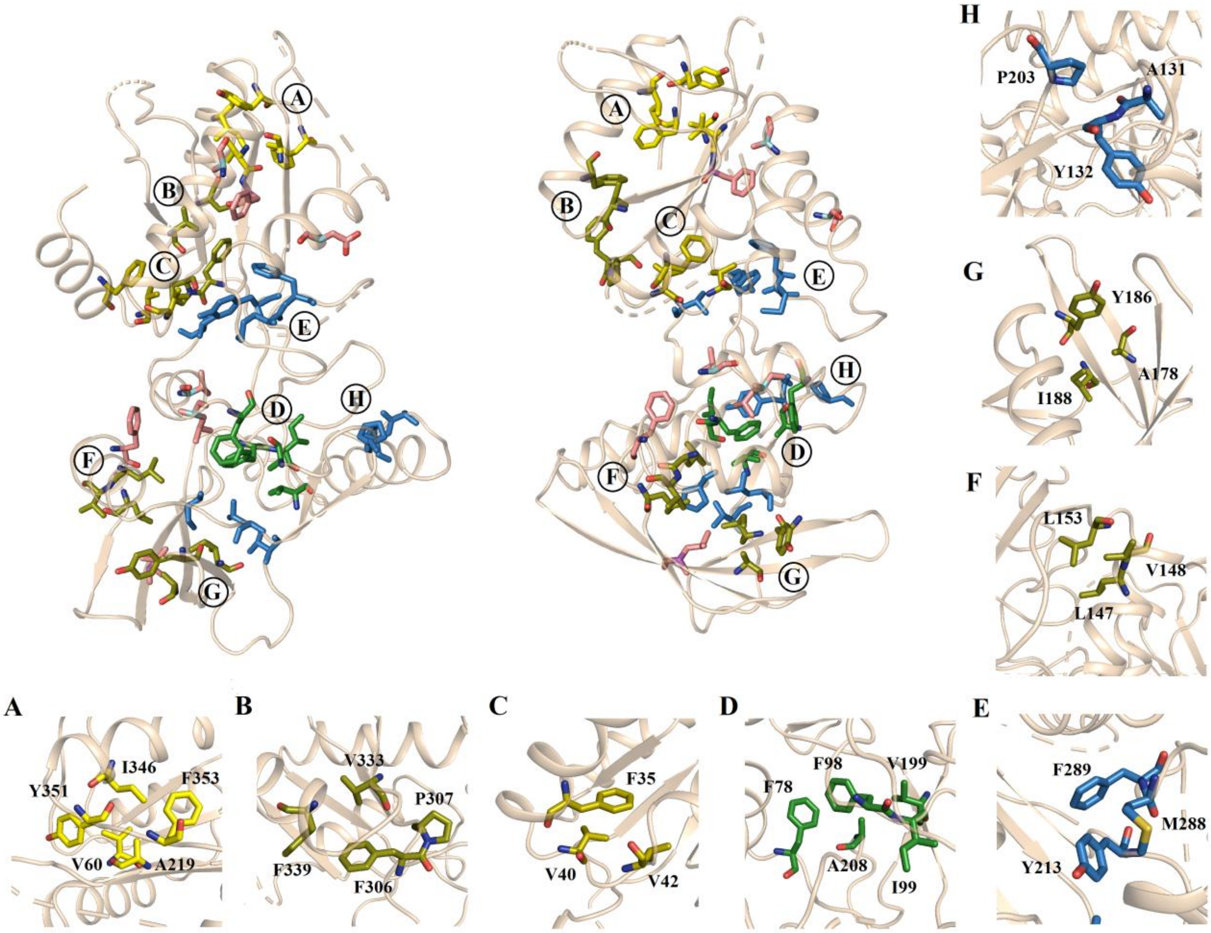


**Figure S9. Hydrophobic clustering in the crystal structure of ExoY.** A minimum of three hydrophobic residues within a radius of (4-5Å) was considered as a hydrophobic cluster(10–13). The hydrophobic clusters and residues are shown in different color code according to its hydrophobicity strength and its distribution in secondary structure. The hydrophobic clusters comprising of 4 or more hydrophobic residues were considered as hard hydrophobic clusters and hydrophobic clusters comprising of only 3 hydrophobic residues were considered as soft hydrophobic clusters. The hard hydrophobic clusters in ordered secondary structures are coded in yellow color whereas in disorder structures are coded in green color. The soft hydrophobic cluster in the ordered secondary structure is represented by dark olive color and in semi-ordered structures such as helix-turn-helix is represented by olive color. The isolated hydrophobic residues in order structures are shown in pink color whereas in disorder structures represented in blue color. A total of 2 hard, 6 soft hydrophobic clusters were found in three- dimensional structure of ExoY. 1 hard (cluster A) and 3 soft (cluster B, G, and F) hydrophobic clusters are found in order structural modules. 1 soft hydrophobic cluster (cluster C) is found in helix-turn-helix. 1 hard hydrophobic cluster (cluster D) and 2 soft hydrophobic clusters (cluster E and H) have simultaneously evolved in disorder globule.

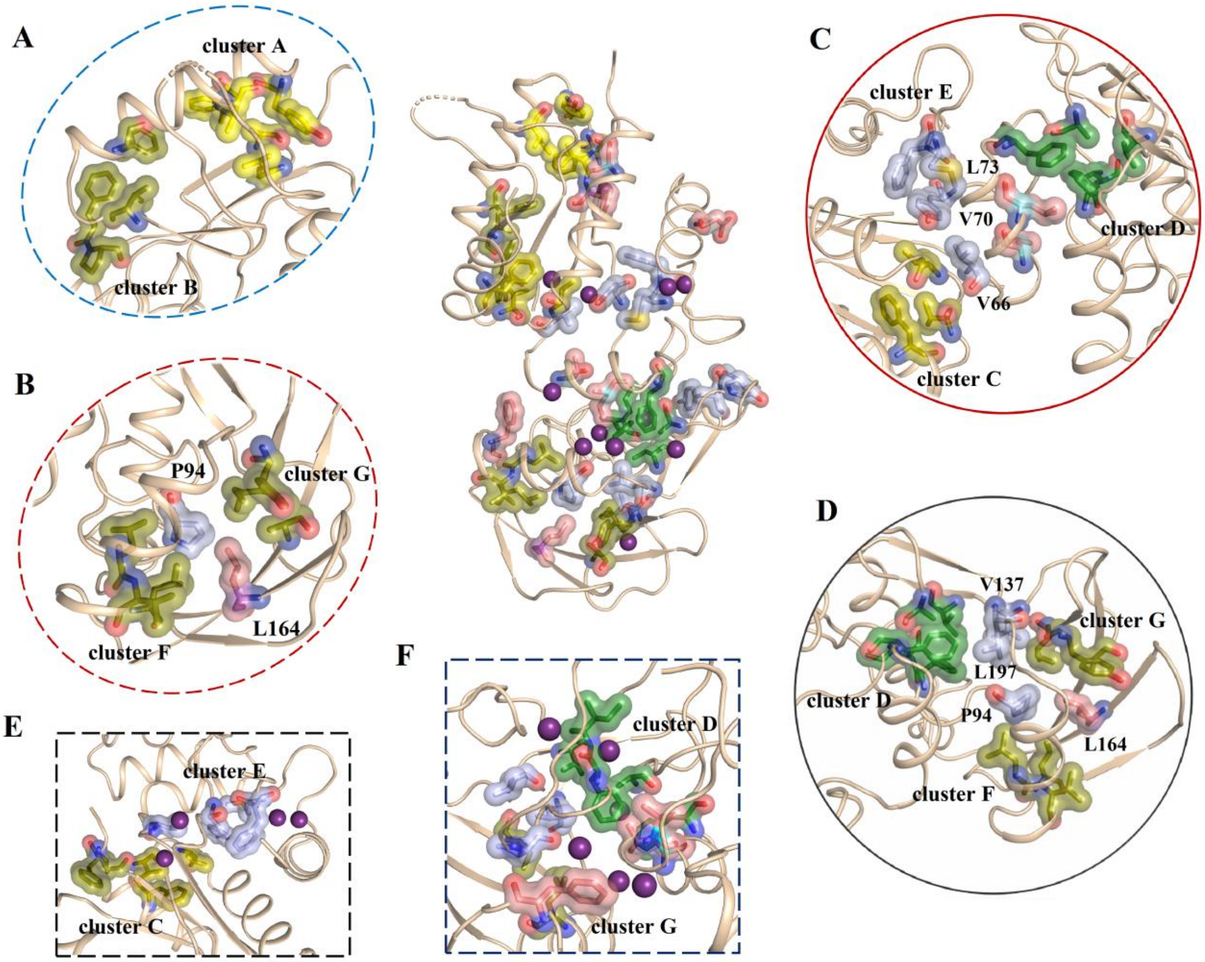


**Figure S10. Hydrophobic packing and geometrical frustration in the folded cores of ExoY.** Cartoon representation of ExoY structure with hydrophobic residues shown in the surface view. The hydrophobic clusters and residues are shown in different color code according to its hydrophobicity strength and its distribution in structure (for more details see **Fig. S9**). (**A**) In ExoY subdomain 2, four beta strands (β1, β8, β9, β10) and one helix (α6) organize to form barrel like structure (barrel 2). The hydrophobic cluster A and cluster B form lids of this empty barrel (portrayed inside black oval shape). (**B**) Similarly, in ExoY subdomain 1, four beta strands (β3, β4, β5, β6) and a helix (α5) also assemble to form twisted barrel like structural fold with hydrophobic cluster F and G acting as its lids (portrayed inside red oval shape). Unlike, barrel structure in lobe 2, it has two isolated hydrophobic residues (P94 and L164) forming weak connections between its hydrophobic lids. The remaining hydrophobic clusters are located in either semi-ordered or disordered structural regions. (**C**) The isolated hydrophobic residues (V66, V70 and L73) form a weak hydrophobic zipping between cluster C, cluster D and cluster E. (**D**) Similarly isolated hydrophobic residues (L137 and V197) form weak hydrophobic connection between hard hydrophobic clusters D in disordered globule with hydrophobic cluster G of barrel like fold in lobe 1. More, upon inspection of water solvation around these hydrophobic clusters, multiple water molecules were found entrapped between hydrophobic clusters and its connecting residues. (**E**) Two water molecules are found between soft hydrophobic cluster C and cluster E. (**F**) Similarly, three water molecules were found to be entrapped between hard hydrophobic cluster D and cluster G of the barrel-like fold. In addition, many more water molecules present around these hydrophobic clusters. Presence of entropically unstable water molecules between hydrophobic clusters and connecting hydrophobic residues further destabilizes the weak hydrophobic interactions. Moreover, it was observed that two water molecules were also found between hydrophobic residues of hard hydrophobic clusters D. Such penetrated water molecules inside hydrophobic cluster increase decrease entropy of water that will subsequently reduce the overall stability of this hydrophobic cluster(10, 11).

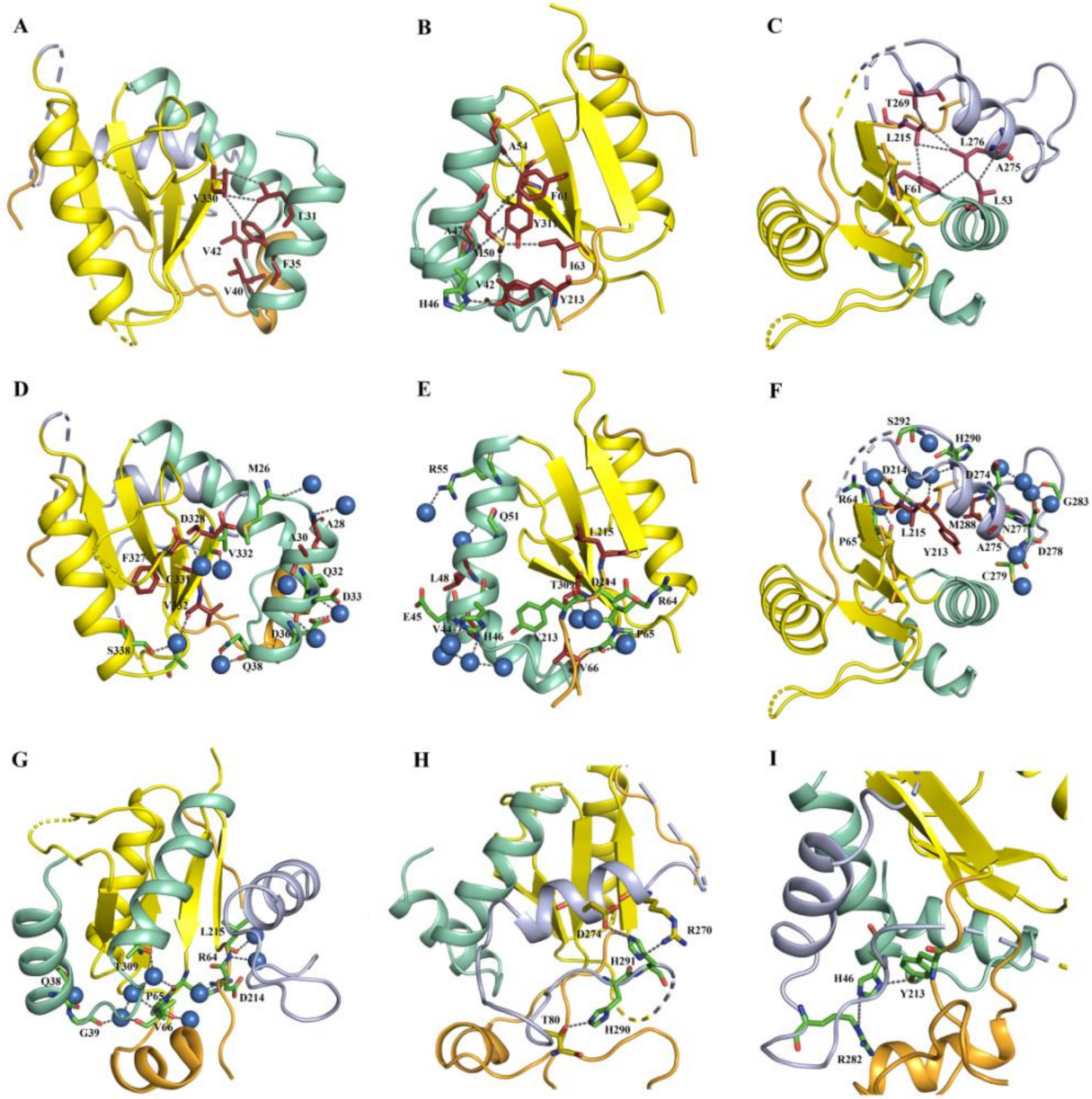


**Figure S11. Decoding the structural stability folded sub-domain 2: Envisaging the non-polar side chain contacts and ion pair interactions with the relatively most stable fold (barrel 2) with interfacial helices.** The sub-domain 2 of ExoY comprised of the stable folded barrel 2 (yellow) linked with N-terminal helices (α1 and α2, green cyan) and C-terminal helix (α7, light blue) via hydrophobic interactions. **(A)** The helix α1 with secretion signal was zipped to barrel structure via three hydrophobic residues (L31, F35, and V330). Interestingly, only one single residue (V330) alone from barrel structure remained critical for a stable interaction between α1 helix and barrel structure. **(B)** Compared to the α1 helix, the number of interaction between α2 helix and barrel structure was relatively higher. The three hydrophobic residues (A47, A54, and M50) from α2 helix interacted with other three hydrophobic residues (F61, I63, and Y211) from barrel structure. The hydrophobic residue (M50) also interacted with hydrophobic residue (V42) of the loop between α1 and α2 helices. Though the hydrophobic contacts were more compare to helix 1, it suffered from destabilization of hydrophobic core partially due presence of amphipathic residues like Y211 and M50. Finally, the interaction between α2 helix and barrel structure was also stabilized by electrostatic interaction between histidine (H46) of the α2 helix with terminal tyrosine residue (Y213) of barrel structure, which might be destabilized during protonation of H46 at a pH below its pKa. **(C)** The C-terminus α7 helix was linked to barrel structure with aid of three crucial residues (L276, A275, and T269) of α7 helix. The residues L276 and T269 interacted directly with hydrophobic residues (F61, L215) of barrel structure. The hydrophobic residues (L276, A275) also interacted with hydrophobic residue (L53) of α2 helix, stabilizing the interaction of α2 and α7 helices with barrel structure. Though the hydrophobic packing of the α7 helix with the folded barrel 1 was most compact, but it was connected with C-terminal disordered flanked of ExoY and suffered from lots of internal side and backbone salvation (discussed later). **Interfacial hydration between barrel 2 and (α1, α2, and α7)**: **(D)** The number of water molecules within 4-5 Å of α1 helix and binding cavities of the α1 helix with barrel structure. A total of 5 water molecules directly formed hydrogen bonding with α1 helix and remaining 4 water molecules were found in the close milieu of binding cavities of α1 helix and barrel structure **(E)** The α2 helix was also formed direct association with 6 water molecules and many more water molecules in the close milieu. **(F)** The level of hydration of α7 helix was very high, with a density of 12 water molecules within 4-5 Å. **(G)** The number of water molecules interacting with barrel structure along the binding region of barrel structure with helices (α1, α2, and α7). **Histidine mediated salt-bridge formation in the interfacial space of between barrel 2 and (α1, α2 and α7): (H)** The histidine residues (H290 and H291) from preceding loop of helix α7 formed interaction with residue (T80) of disorder globule of ExoY and with residues (R270 and D274) of α7 helix. These interactions are crucial for linking folded domain 2 with disorder globule of ExoY that subsequently contributes to overall structural stability. **(I)** Another histidine residue (H46) of α2 helix simultaneously formed interaction with terminal tyrosine residue (Y213) of the barrel and with a terminal arginine residue (R282) of the loop associated with the α7 helix. The PMF mediated lowering of pH below the pKa of histidine might possibly disrupt these crucial interactions mediated by histidine residues. This might subsequently increase water penetration inside the interfacial binding cavity; thereby destabilizing the interactions between barrel 2 and helices (α1, α2 and α7) during pH-induced partial unfolding and might sometimes cause unfolding of helix conformers as observed from our experimental studies (Raman crystallography and FT-IR analysis) of secondary structure unfolding and tertiary structural hydration.

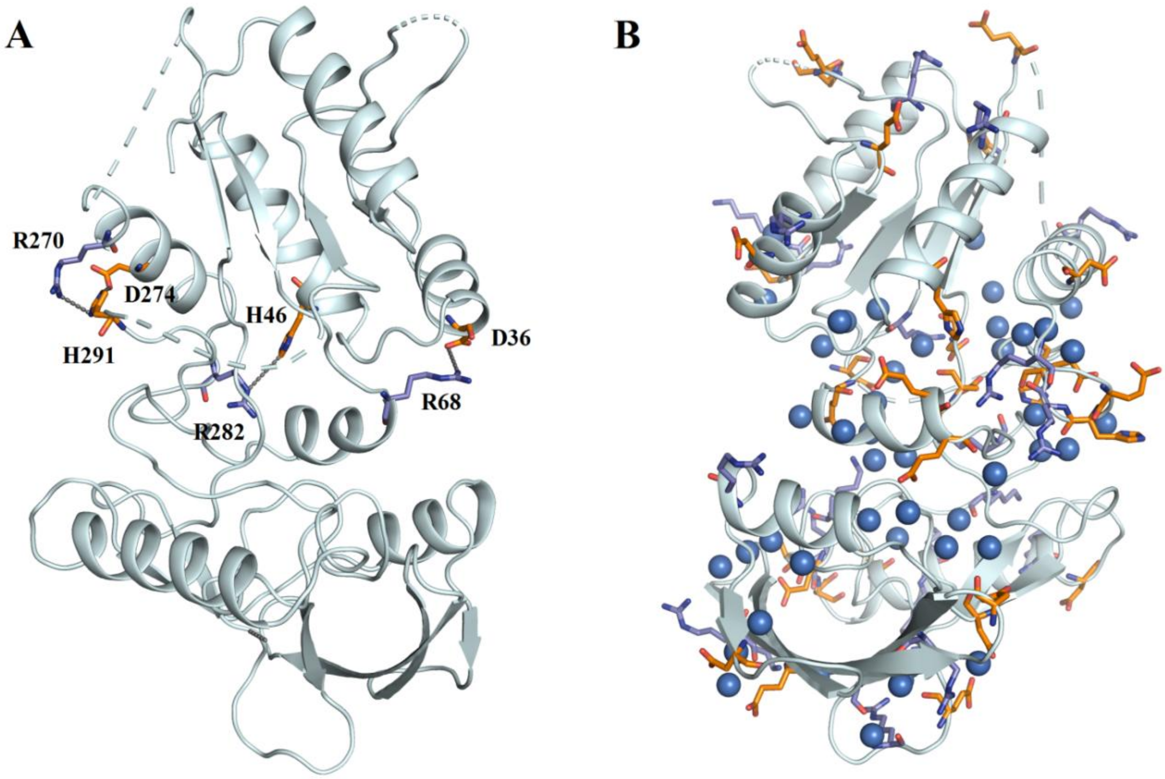


**Figure S12. Distribution of charged residues, and long-range electrostatic interactions between oppositely charged residues in ExoY. (A) Restricted residue-residue interaction:** Among 34 positively (Arg and Lys) and 33 negatively (Asp and Glu) charged residues in the crystal structure of ExoY, only 7 residues were found to involve in long-range electrostatic ion-pair interactions [(R270, D274 & H291), (D36 & R68), and (H46 & R282)]. **(B) Labile tertiary structure:** The majority of charged residues formed extensive interaction with bulk water molecules or formed water-mediated dipolar interactions with other polar and charged residues.

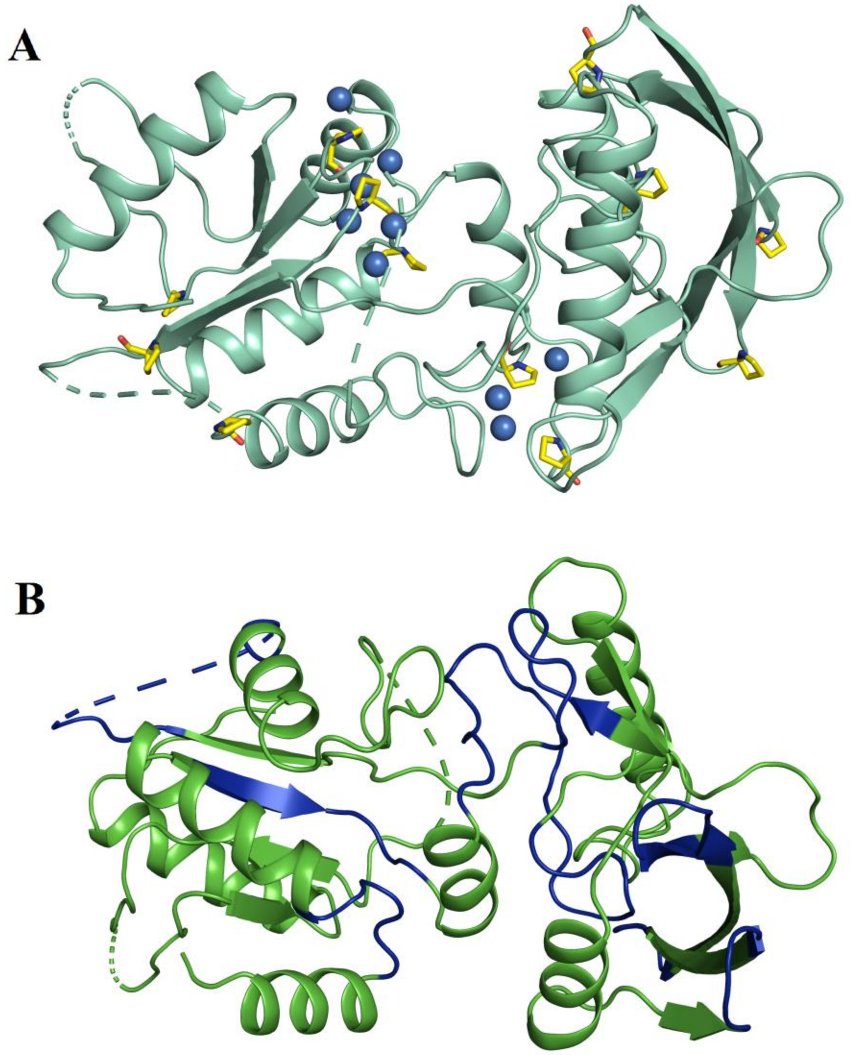


**Figure S13. Evolutionary impact of proline and glycine-proline enriched ELP-like peptide motifs in ExoY. (A)** Proline residues in ExoY are more strategically positioned in the flanking region of helices and β-strand segments (marked in yellow color) and contain internal water molecule (shown in blue color) which may facilitate early stage unfolding events. As proline residues introduce strong selection pressure on folding (helix and β-breaker residue) and the existence of hydrated cis-proline in sequence landscape not only restrain local and global folding (14) (critical for effector unfolding and secretion), but might also serve as a gatekeeper residues which prevent initiation of amyloid-like aggregation (rich in cross-β structures) of intermediately folded unstable effector proteins (7). Consistent with this, rapid formation and stabilization of extended PPII or type II β-turn in early stage unfolding of ExoY was observed in FT-IR and Raman unfolding experiments (**Fig. 2 & S2** and **Table S1 & S2**). **(B)** Abundance of glycine–proline rich elastin-like peptide sequences or motifs (blue) in the folded cores of ExoY. As ELP-like motifs have higher entropic cost for secondary structure formation and proline residues exhibit structural constraints for the formation of β-sheet or cross-β-structure (14–16). Glycine–proline rich ELP-like peptide sequences might also play a significant role in the oil-water-oil type of phase separation. Consistently, even though ExoY exhibits a higher propensity for self-association, still, it does not form typical amyloid-like aggregates (rich in cross-β-structures). Furthermore, the increase in coacervation of the propensity of ExoY in presence of mild acidic conditions of PMF indicate increased hydration of disordered globule (rich in ELP-like sequences) and separation of folded barrel structures (**Fig. S6**). Thus, the positive evolution of proline and LCR promoting residues highlight possible generic strategies or complementary mechanisms in T3SS effector proteins to prevent amyloid-like aggregation and to maintain a fine balance in folding, solubility, and stability (6, 7, 15).

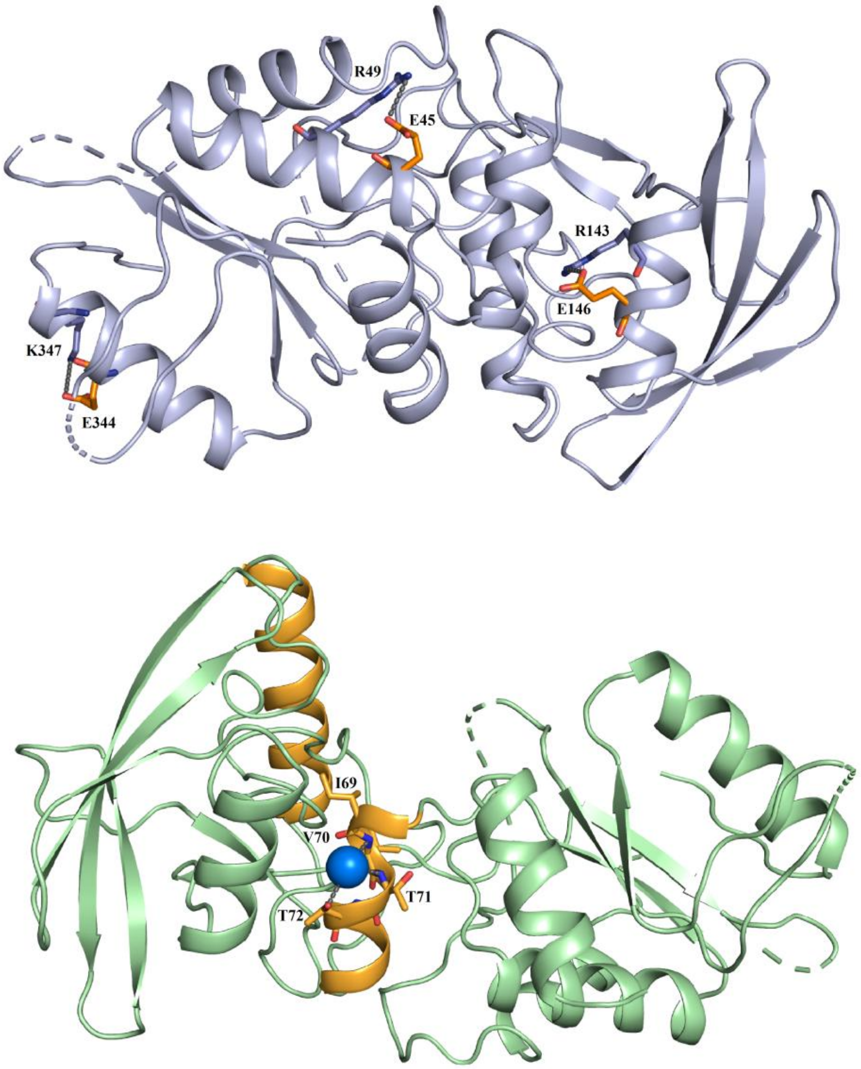


**Figure S14. Potentially optimized helix stabilizing and destabilizing interactions in isolated helices of ExoY. (A)** Additional helix stabilization via helix anchoring salt bridge formation (short-range electrostatic interactions between oppositely charged ion pairs at i and i+4 or i+3 positions) at the terminal regions of three helices (α2, α5 & α7) in ExoY. Since glutamate residues played the role of negatively charged residue in these ion-pairs interactions, a significant loss in secondary structures (helices) observed at extremely low pH < 4 (**Fig. S5**) is possibly due to destabilization of salt bridge interactions (pKa of Asp or Glu). **(B)** The isolated helices (α2, α3; orange color) with no significant interactions or associations with barrel-like structures or folded cores of ExoY. Potential destabilizing role of water molecule encapsulated in a kinked region of isolated helix α3 by preventing the formation of pure α-helix via backbone amide interaction. Furthermore, it also highlights the potential conformational frustrations which facilitate the formation of extended PPII-like helix conformations at lower pH conditions.

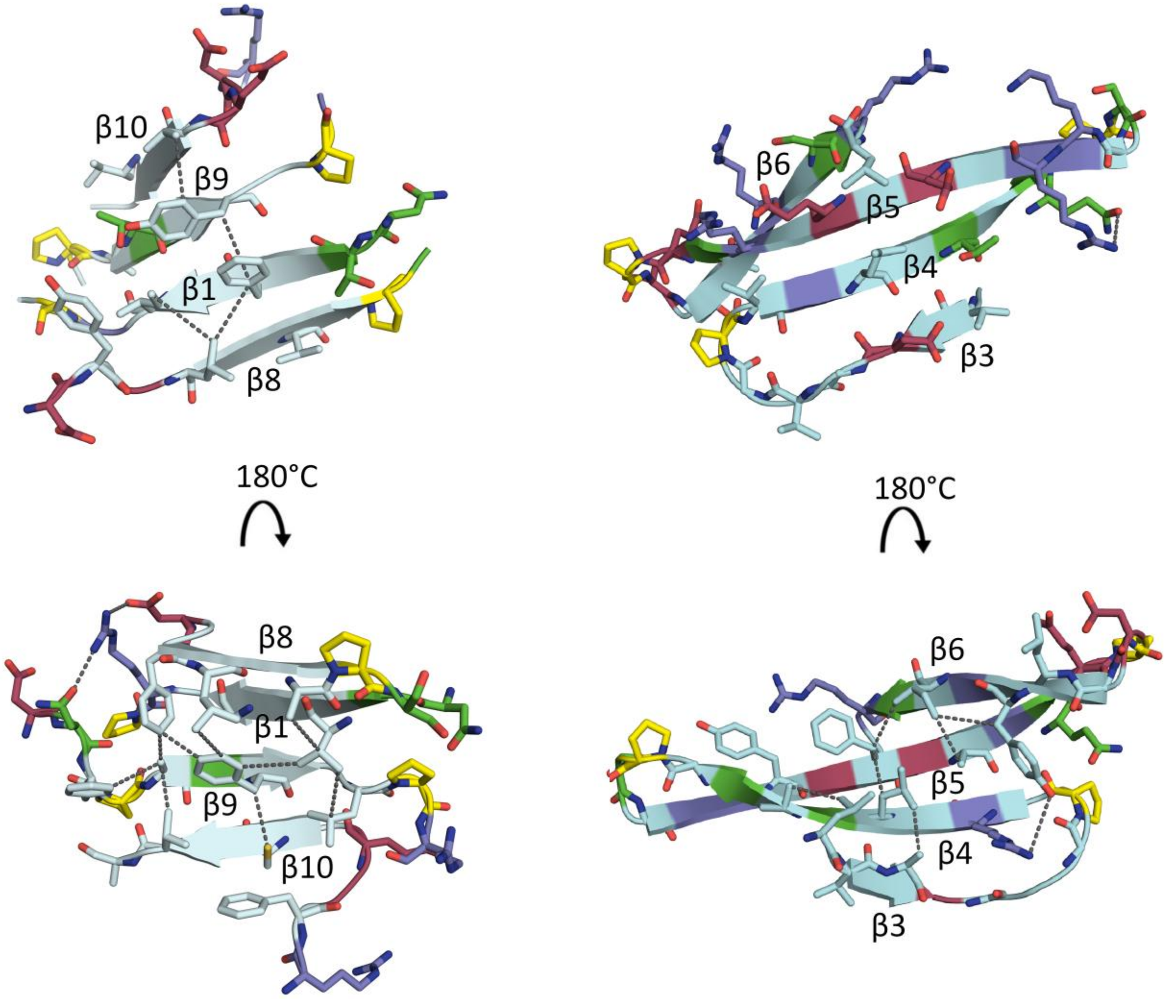


**Figure S15. Geometrical stress in short or amphipathic twisted β-blade conformations embedded in the core of relatively stable barrel-like folds.** ExoY comprises short and amphipathic β-strands that are frequently terminated with β-breaker proline residues (marked in yellow color). More, due to weak hydrophobic contacts and amphipathic nature of side chain residues embedded with each β-strands segment, the stability of β-strands is significantly compromised, and β-strands segments attained twisted β-bladed conformations. Individual β-strands in β-bladed conformations (left panel) are extremely short and frequently terminated by proline residues though there were potential hydrophobic interactions between side chains (marked as dotted lines). In right panel, the individual β-strands in β-bladed conformations are relatively more amphipathic in nature.

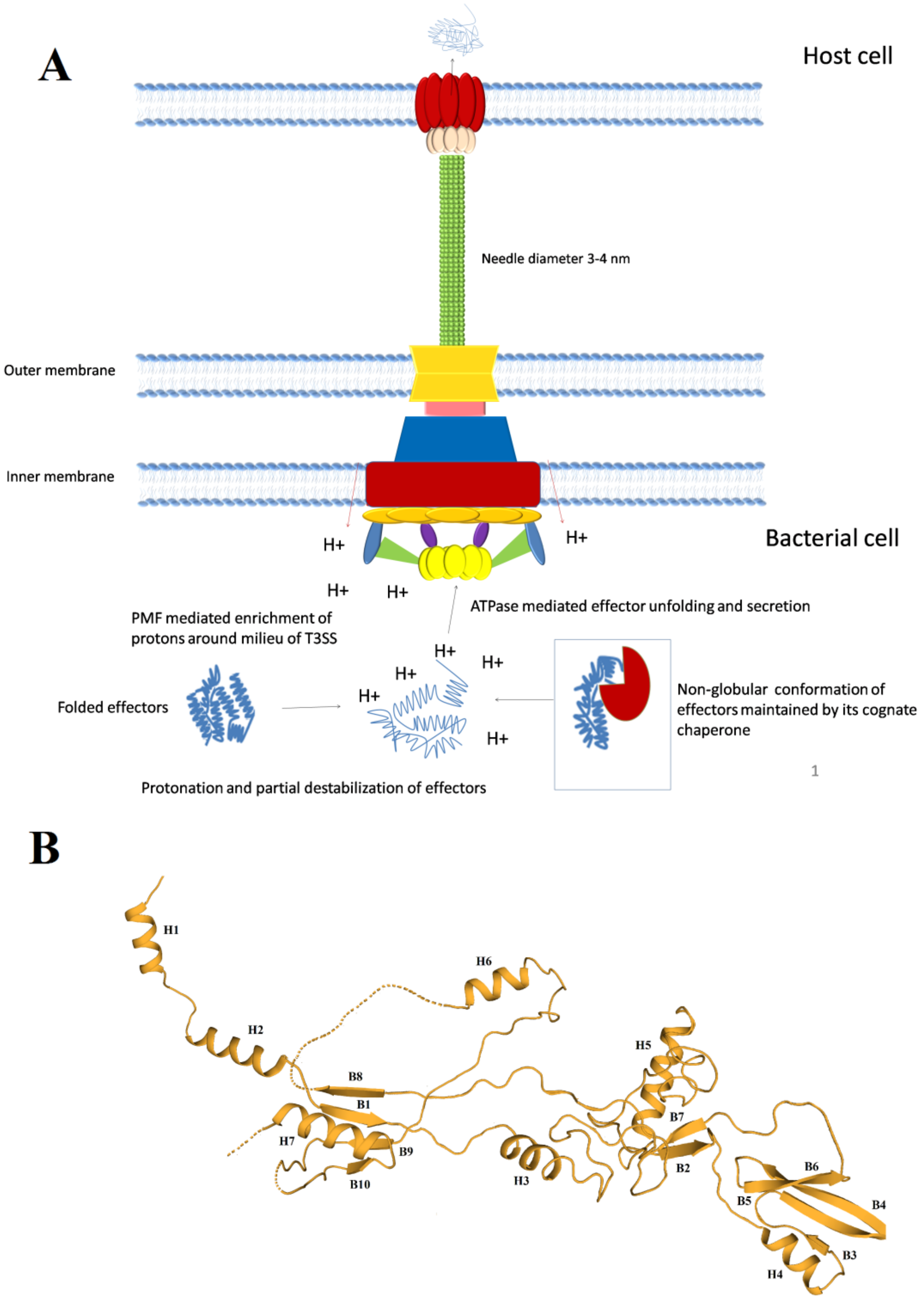


**Figure S16. The hypothetical model for the mechanism of effector protein unfolding and T3SS secretion. (A)** The working model showing the possible steps in effectors' unfolding for T3SS secretion. In the course of evolution, multiple factors that govern the stability have been under strict natural selection in order to reduce the energetic cost of effectors' unfolding. The proton motive force (PMF) which primarily energizes the rate-limiting step of type-III secretion, can also lower pH in the close milieu of T3SS assembly. This phenomenon of lowering of pH in the close milieu of T3SS secretion has been intelligently exploited by reduces the folding and stability of effectors. The pH-dependent unfolding of effectors not only minimizes the energetic cost of complete effectors' unfolding by ATPase but also accelerate the rate-limiting step of effector protein unfolding for T3SS secretion. **(B)** Hypothetical model depicting of PMF-mediated partially unfolded state of ExoY (contains typical block copolymeric geometry with two folded barrels as evaluated from decoding of overall structural stability of ExoY) that might accommodate narrow passage of T3SS apparatus.

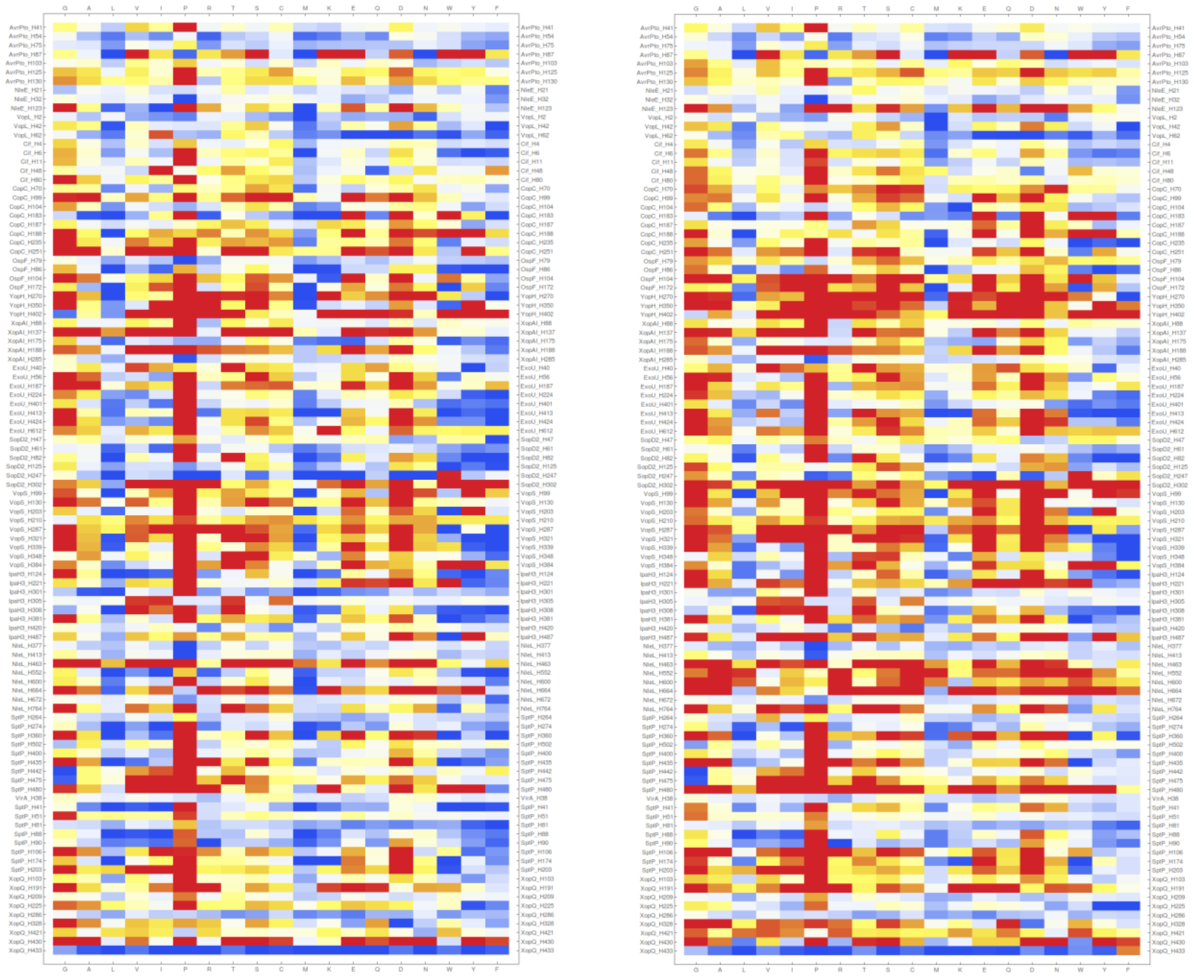

pH 7.4 pH 5.8

**Figure S17. Distribution of stability effects of in-silico mutagenesis of histidine residues in T3SS effectors.** The total of 104 histidine residues from 16 PDB structures from 11 bacterial species was mutated individually into 19 other amino acids. The energetic effect of mutations was classified into different classes; highly destabilizing (red), moderately destabilizing (orange), slightly destabilizing (yellow), neutral (green), slightly stabilizing (blue), moderately stabilizing (indigo), and highly stabilizing (violet). From the in-silico mutagenesis analysis carried out at pH 7.4, it was clearly observed that almost 1/3rd of the histidine residues (33 of 104) had accumulated almost 3/4th of the total highly destabilizing mutations ((232 of 306 = 75.81%), ($\Delta\Delta G_{fold}$ per chain > +1.84 kcal/mol)) and seemed to be critical for the stability of native protein conformations. Further, a significant variation in distributions of stability effects of mutations at pH 7.4 (Right) and pH 5.8 (Left) can be seen, driven by the differential local environment. On average, the distributions of destabilizing mutations was lesser (33.30% vs. 25.81%), and the distributions of stabilizing mutations were more (13.31% versus 18.47%) at pH 5.8 compared to pH 7.4. Increase in stabilizing and neutral mutations at low pH revealed the destabilizing nature of histidine residues at low pH.

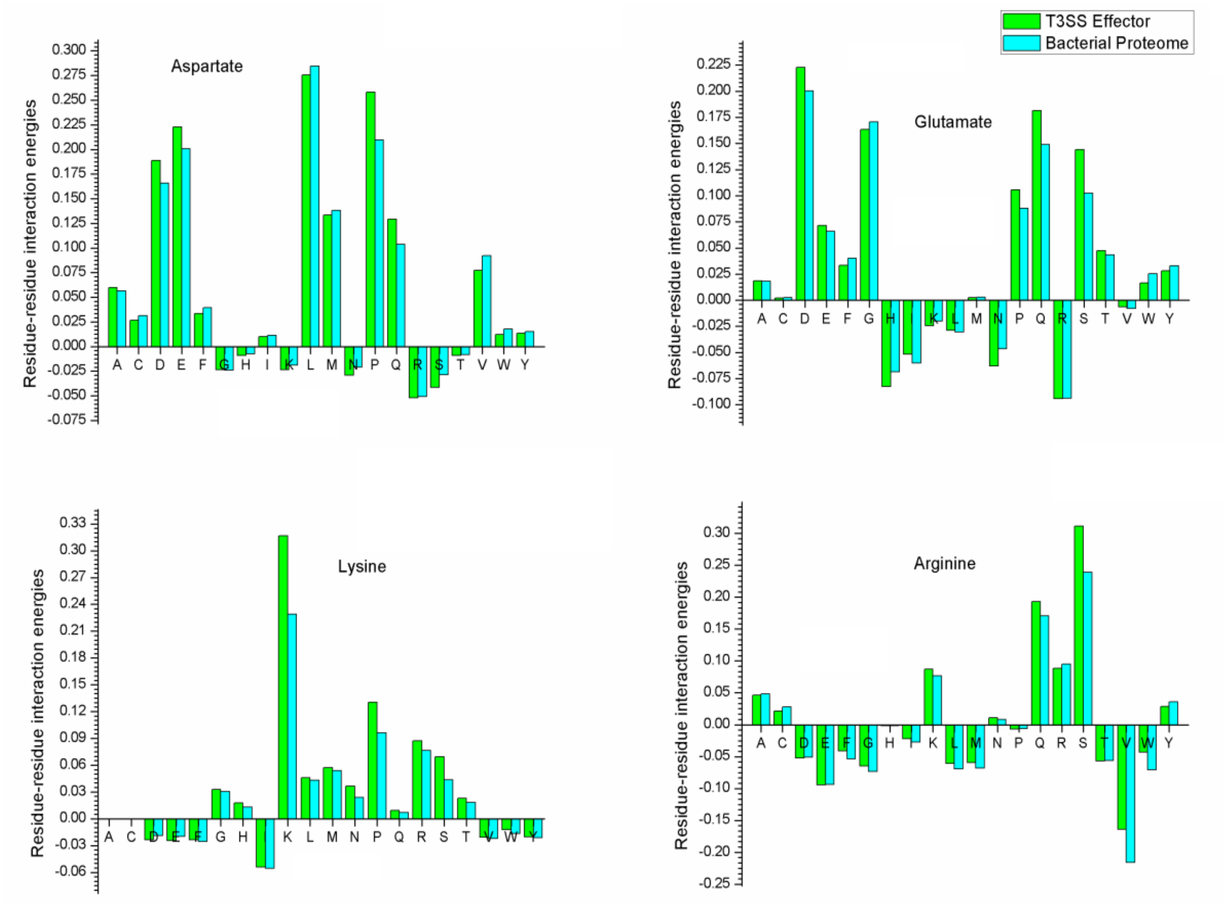


**Figure S18A.** A detailed depiction of pairwise residue-residue interaction energies of charged amino acids (Asp, Glu, Lys, and Arg) with all 20 naturally evolved amino acids in T3SS effectors and control bacterial proteome respectively.

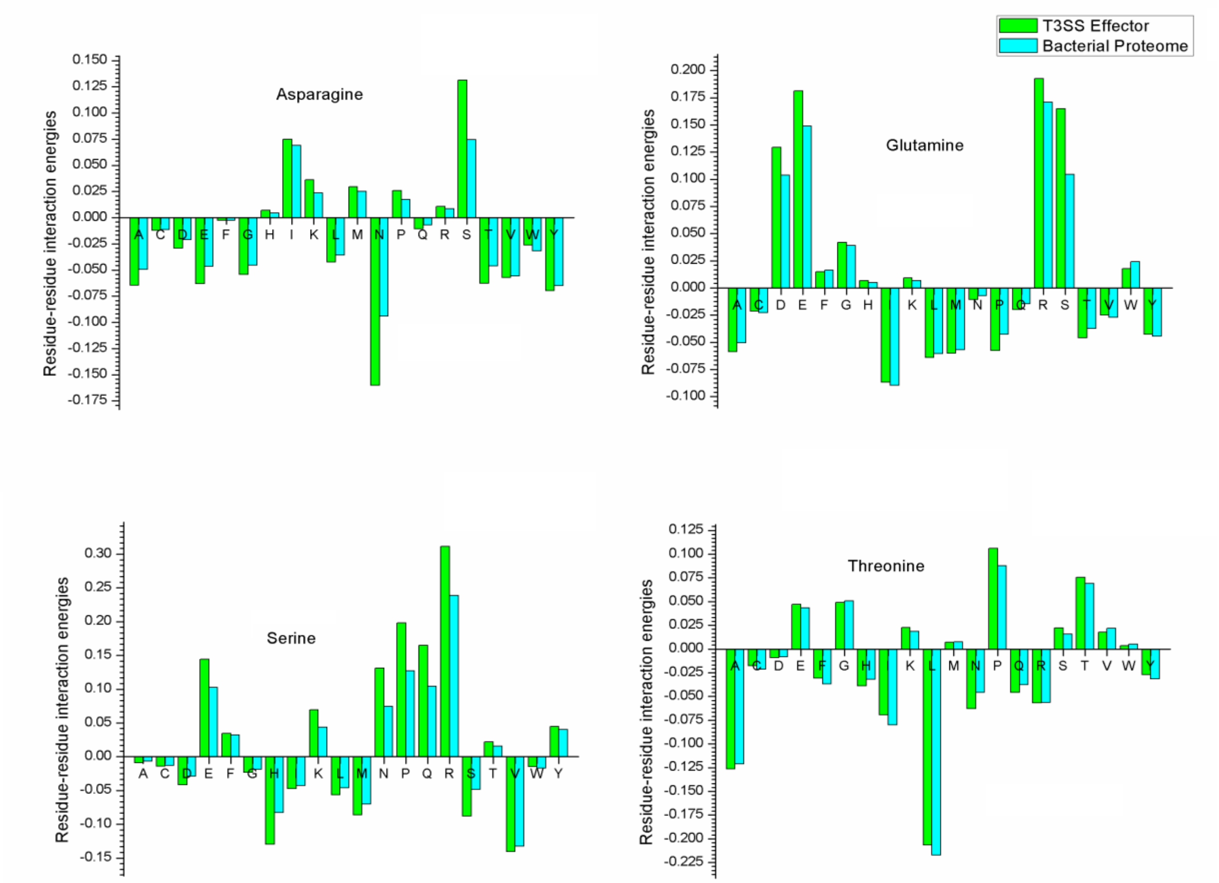


**Figure S18B.** A detailed depiction of pairwise residue-residue interaction energies of polar amino acids (Asn, Gln, Ser, and Thr) with all 20 naturally evolved amino acids in T3SS effectors and control bacterial proteome respectively.

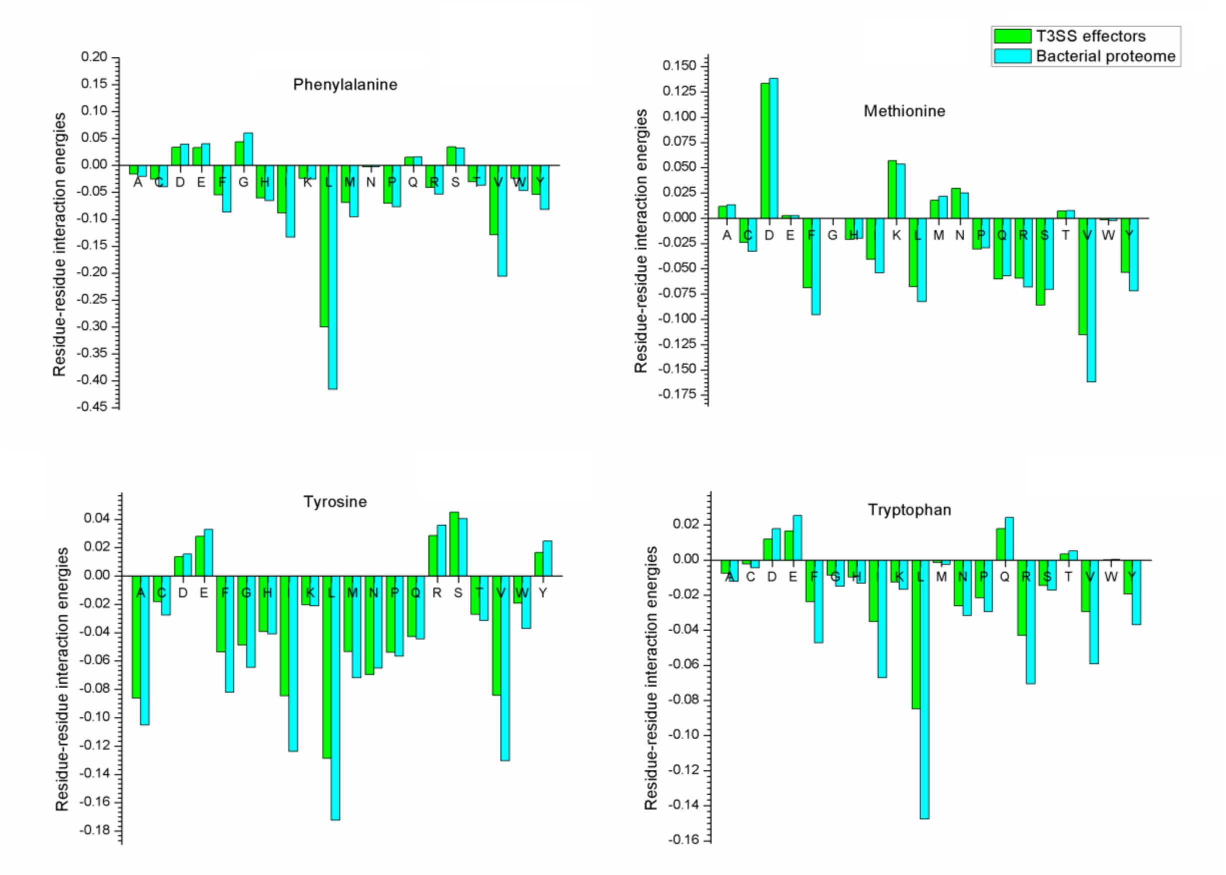


**Figure S18C.** A detailed depiction of pairwise residue-residue interaction energies of hydrophobic aromatic amino acids (Phe, Tyr, and Trp) and methionine with all 20 naturally evolved amino acids in T3SS effectors and control bacterial proteome respectively.

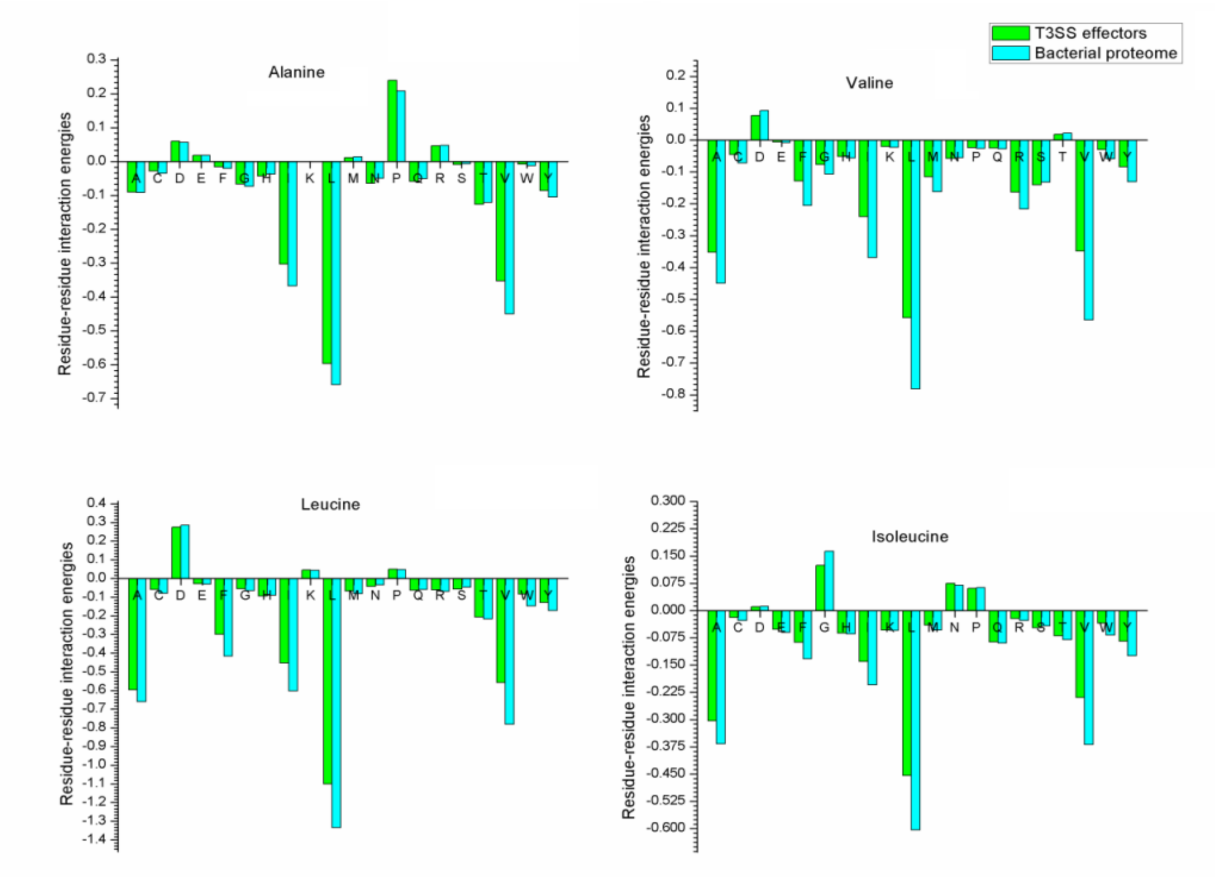


**Figure S18D.** A detailed depiction of pairwise residue-residue interaction energies of hydrophobic non-aromatic amino acids (Ala, Val, Leu, and Ile) with all 20 naturally evolved amino acids in T3SS effectors and control bacterial proteome respectively.

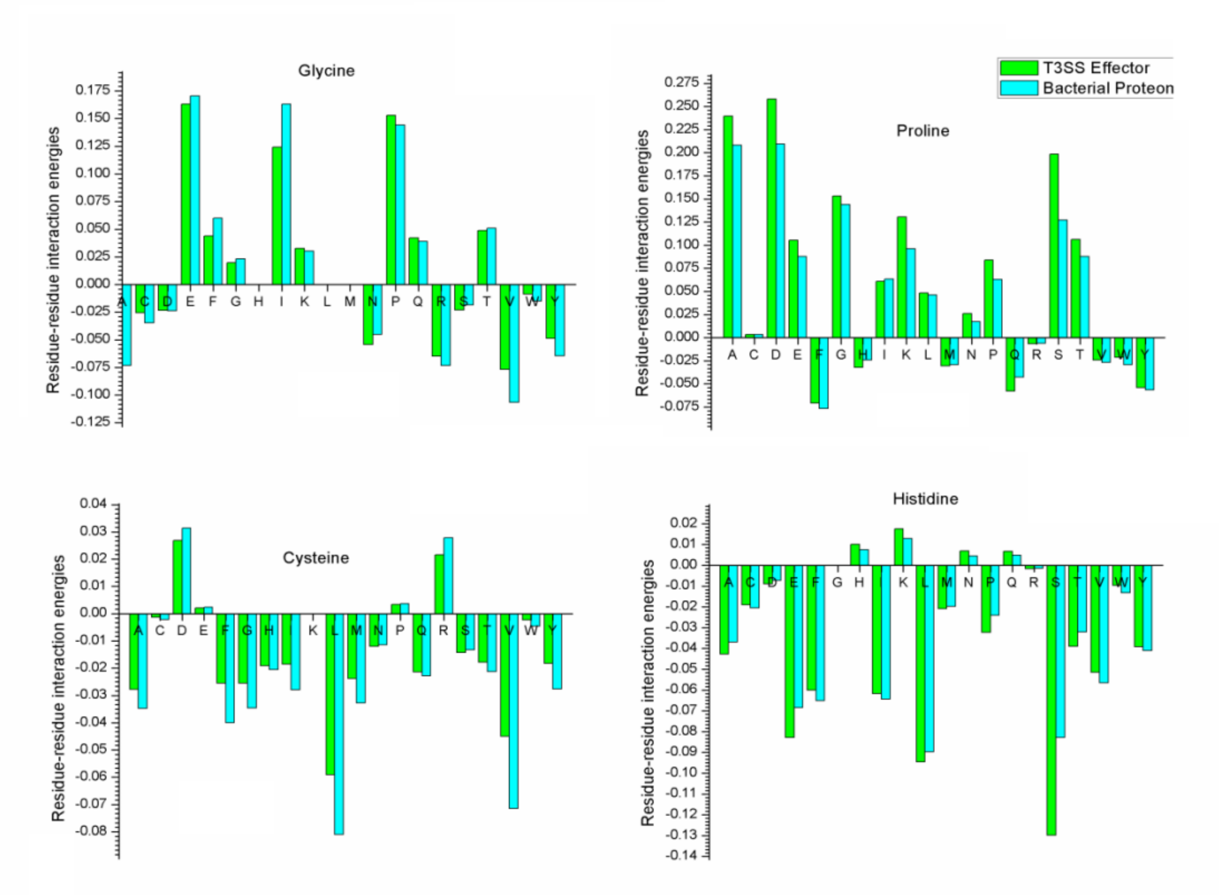


**Figure S18E.** A detailed depiction of pairwise residue-residue interaction energies of glycine, proline, cysteine and histidine residues with all 20 naturally evolved amino acids in T3SS effectors and control bacterial proteome respectively.
